# Separable Spatial Single-cell Transcriptome Representation Learning via Graph Transformer and Hyperspherical Prototype Clustering

**DOI:** 10.1101/2025.05.01.651634

**Authors:** Laiyi Fu, Penglei Wang, Gaoyuan Xu, Jitao Lu, Hequan Sun, Danyang Wu

## Abstract

Spatial transcriptomics enables the exploration of cell types, spatial domain organization, and cell–cell communication at tissue scale, serving as a powerful link between tissue morphology and molecular function. However, existing methods still struggle with limited spatial resolution, poor robustness in multi-slice alignment, and suboptimal ligand-receptor (L-R) detection in complex tissues. To address these challenges, we propose S3RL (Separable Spatial Single-cell transcriptome Representation Learning via Graph Transformer and Hyperspherical Prototype Clustering), a unified framework that integrates gene expression, spatial coordinates, and histological image features via a graph neural network and hyperspherical prototype-based separable representation learning. In spatial clustering tasks, S3RL improves the average Adjusted Rand Index (ARI) by nearly 120% on the Nanostring lung cancer dataset and over 26% on the 10X DLPFC dataset compared to state-of-the-art methods. For multi-slice spatial alignment, S3RL achieves an average ARI improvement of over 65.4% in partial brain slices and over 48.3% in complete slices relative to GraphST, highlighting its superior robustness and alignment consistency across heterogeneous spatial inputs. Moreover, S3RL reveals more biologically meaningful ligand-receptor signaling relationships, enhancing the interpretability of cell-cell communication patterns. Together, these results demonstrate S3RL’s effectiveness in enhancing spatial resolution, cell type identification, and biological insight across diverse spatial transcriptomics datasets.

## I. Introduction

In multicellular tissues or organisms, cells with similar biological functions tend to organize into coherent clusters, forming spatially structured domains that are essential for maintaining physiological function and developmental coordination. These spatial arrangements underlie the architecture of diverse organs—such as layered neuronal structures in the brain, glandular compartments in epithelial tissues, and immune cell niches in lymphoid organs—and are key to understanding how local cellular interactions contribute to global tissue behavior. Disruptions to these spatial organizations are often associated with pathological conditions, including cancer, neurodegeneration, and inflammatory diseases. Therefore, accurately capturing the spatial layout of functionally related cells is crucial for studying tissue organization, identifying disease-related microenvironments, and elucidating the spatial dynamics of intercellular communication networks.

To address the need for spatially resolved molecular insights, the field of single-cell spatial transcriptomics has witnessed remarkable advancements in recent years [1]. While conventional single-cell RNA sequencing (scRNA-seq) technologies have proven highly effective in dissecting cellular heterogeneity, they inherently lack spatial information, which is indispensable for interpreting tissue structure and intercellular communication. To overcome these limitations, a variety of spatial transcriptomics techniques have been developed that aim to jointly preserve spatial context and capture transcriptomic profiles at high resolution. However, reconstructing spatial information at single-cell resolution with exhaustive gene coverage remains challenging due to technical variability, gene dropout, and high data sparsity [2]. For example, 10X Visium [3] captures transcriptomic signals from spots that often span multiple cells, limiting its ability to resolve fine-scale features. Although 10X Visium HD offers improved resolution, it suffers from shallow sequencing depth and increased cost. Emerging technologies such as Slide-seq [4], [5], DBiT-seq [6], and Stereo-seq [7] provide enhanced spatial granularity but still face challenges such as limited sensitivity and gene coverage [8]. Likewise, FISH-based methods like SeqFISH [9] deliver multiplexed spatial data but are constrained by the number of genes that can be simultaneously profiled.

Recent advances in spatial transcriptomics technologies have enabled the simultaneous generation of multimodal data, including gene expression profiles, spatial coordinates, and histological images (such as H&Estained sections) [3], [4], [7]. These sequencing platforms themselves offer the technical foundation for collecting such complementary data types. The integration of gene expression with tissue morphology provides new opportunities to analyze cell function and organization in situ, especially in spatially heterogeneous tissues [8]. By mapping gene expression patterns to distinct spatial niches and cellular phenotypes, spatial transcriptomics analysis can reveal novel biological insights into processes like cell fate determination, microenvironmental regulation, and cell-cell communication networks [10]– [12].

However, effectively leveraging this multimodal information requires downstream computational strategies that can model and fuse these diverse data sources. Various approaches have been proposed to achieve this goal. For example, Giotto [13] employs Hidden Markov Random Fields to capture spatial continuity, while SpaGCN [14] and CCST [15] use graph convolutional networks to incorporate spatial and gene expression features. StLearn [16] extracts morphological features from histological images and integrates them with transcriptomic data. Bayesian methods such as BayesSpace [17] improve spatial resolution using probabilistic inference. More recently, models like STAGATE [18], GraphST [19], SiGra [20], and SEDR [21] utilize graph autoencoder-based frameworks to learn latent spatial representations. In general, while the sequencing platforms generate rich multimodal datasets, it is the computational models that play a central role in integrating and interpreting this information. The synergy between advanced data acquisition technologies and increasingly sophisticated analysis frameworks continues to push the frontier of spatial transcriptomics.

Despite the growing ability of spatial transcriptomics (ST) technologies to integrate multimodal data, several critical challenges remain unaddressed in existing analysis frameworks. First, histological images—despite being a unique modality in ST—often exhibit high structural homogeneity, blurred boundaries, and subtle color variation, which makes it difficult for conventional models to extract meaningful semantic features. For instance, models like SiGra [20] flatten the three channels of H&E-stained images and input them into neural networks. However, due to the aforementioned issues in histological images, such approaches fail to effectively distinguish adjacent regions with distinct biological functions. This highlights the broader limitation of relying on low-level image features, which often contribute little to meaningful spatial representation. Consequently, the utility of image information in multimodal fusion remains limited. Second, most spatial graph construction strategies rely solely on spatial proximity (e.g., Euclidean distance or k-nearest neighbors) such as [14], [19], [21] etc., ignoring functionally similar but spatially distant spots and often introducing noisy connections between spatially adjacent yet biologically distinct regions, limiting the biological relevance of the graph structure. Third, during representation learning and clustering, current methods typically lack constraints on cluster separability in the latent space. This results in cluster overlap, poor inter-cluster separation, and unbalanced clustering outcomes, which in turn limits biological interpretability.

To address these challenges, we propose a framework named Separable Spatial Single-cell Transcriptome Representation Learning via Graph Transformer and Hyper-spherical Prototype Clustering, termed as S3RL. Our method integrates spatial location, histological images, and gene expression within a unified graph-based model. It first extracts high-level semantic features from histology images using contrastive learning, and combines them with gene expression similarity to construct a spatial graph with positive and negative edges—where positive edges reflect potential homogeneity and negative edges encode functional heterogeneity. This dualmodality-guided graph better captures both spatial relationships and biological function connections. On top of this, S3RL learns low-dimensional embeddings constrained on a unit hypersphere with evenly distributed prototypes. A hyperspherical regularization, which is imposed to encourage spots of different types to be uniformly distributed across the hyperspherical manifold, promotes cluster separability and balance, while a pseudo-label-driven prototype update mechanism dynamically adapts to dataset-specific or histology-specific variation, ultimately enhancing both clustering accuracy and generalization. S3RL allows for end-to-end learning of cell type clusters and ensures that learned separable representations capture both spatial context and cellular heterogeneity. We benchmarked S3RL across a wide range of spatial transcriptomic datasets, including 10X DLPFC, Nanostring lung cancer, human breast cancer, mouse brain, Stereo-seq, and Slide-seqV2. On the Nanostring lung cancer dataset, S3RL achieves an average ARI improvement of nearly 120% over all baseline methods, and a 52% improvement compared to the second-best method. On the 10X DLPFC dataset, it outperforms competing models by over 26% relative to the second-best baseline. Furthermore, S3RL demonstrates strong robustness in cross-slice alignment, with up to 27% improvement even under partially missing slice conditions. In ligand-receptor analysis, S3RL uncovers clearer and biologically specific signaling interactions, further supporting its relevance for downstream interpretation tasks.

## II. Results

### A. Overview of the S3RL method

The S3RL model comprises three key modules: the semantic information extraction module, the graph construction module, and the separable representation learning module. In the semantic information extraction module, S3RL adopts the SimCLR framework [22] to extract high-level semantic information from histological images. In the graph construction module, S3RL first constructs the *k*-nearest neighbor graph based on the relative locations from spatial transcriptomics data. S3RL further integrates positive (negative) pairs with high (low) semantic similarity and gene expression similarity into the graph structure to detect non-neighboring similar spots and separating neighboring spots of different cell types. S3RL model leverages the powerful graph neural networks (GNNs) [23] as the backbone of the separable representation learning module and learns the low-dimensional expressive representation of spots by collaboratively taking into account the spatial information (locations of spots), the semantic information (histological images of spots), and the gene expression profiles of spots in the spatial transcriptomics context. Subsequently, S3RL retrieves clustering results in an end-to-end manner via a hypersphere prototype learning submodule. Specifically, the representations are mapped and constrained to the unit hypersphere with evenly distributed prototypes, where each prototype represent the center of a specific cell type. In this way, the affinity of each spot to the prototypes corresponding to different cell types is measured by the cosine similarity between the representations of the spot and prototypes. S3RL further introduces a hyperspherical regularization to ensure that all spots are evenly and closely distributed across prototypes and thus promote separability of the learned representations. Additionally, the current clustering results acting as pseudo labels provide a supervision signal to update the prototypes dynamically, which facilitates the adaptation of the prototypes to histological variations and thus improves the clustering performance.

### B. S3RL’s spatial clustering of human dorsolateral prefrontal cortex 10x Visium data improves layer identification

To evaluate the spatial clustering performance of S3RL on DLPFC tissue slices, we conducted an extensive comparison using the 10X Visium dataset [24]. This dataset includes spatially resolved transcriptomic profiles from 12 DLPFC slices, depicting four or six cortical layers along with the white matter (WM) region. It consists of gene expression data for 33,538 genes collected from three independent neurotypical adult donors, where each set contains two adjacent 10 *µ*m tissue sections, with the second set positioned 300 *µ*m posterior to the first.

We compared S3RL against BayesSpace [17], Giotto [13], Seurat [25], SiGra [20], conST [26], SpaceFlow [27], spaGCN [14], STAGATE [18], SEDR [21], and GraphST [19], using manually annotated cortical layers as ground truth (Figure 2A-C, detailed clustering results in Supplementary Fig. S1 and Fig. S2). S3RL achieved the highest mean clustering accuracy with an ARI of 0.65, outperforming the average of all other methods by 26.8%, and surpassing the best-performing baseline (SiGra) by 10.2%. It demonstrated superior ability in preserving layer boundaries and minimizing over-segmentation(Figure 2D).

**Fig. 1.**
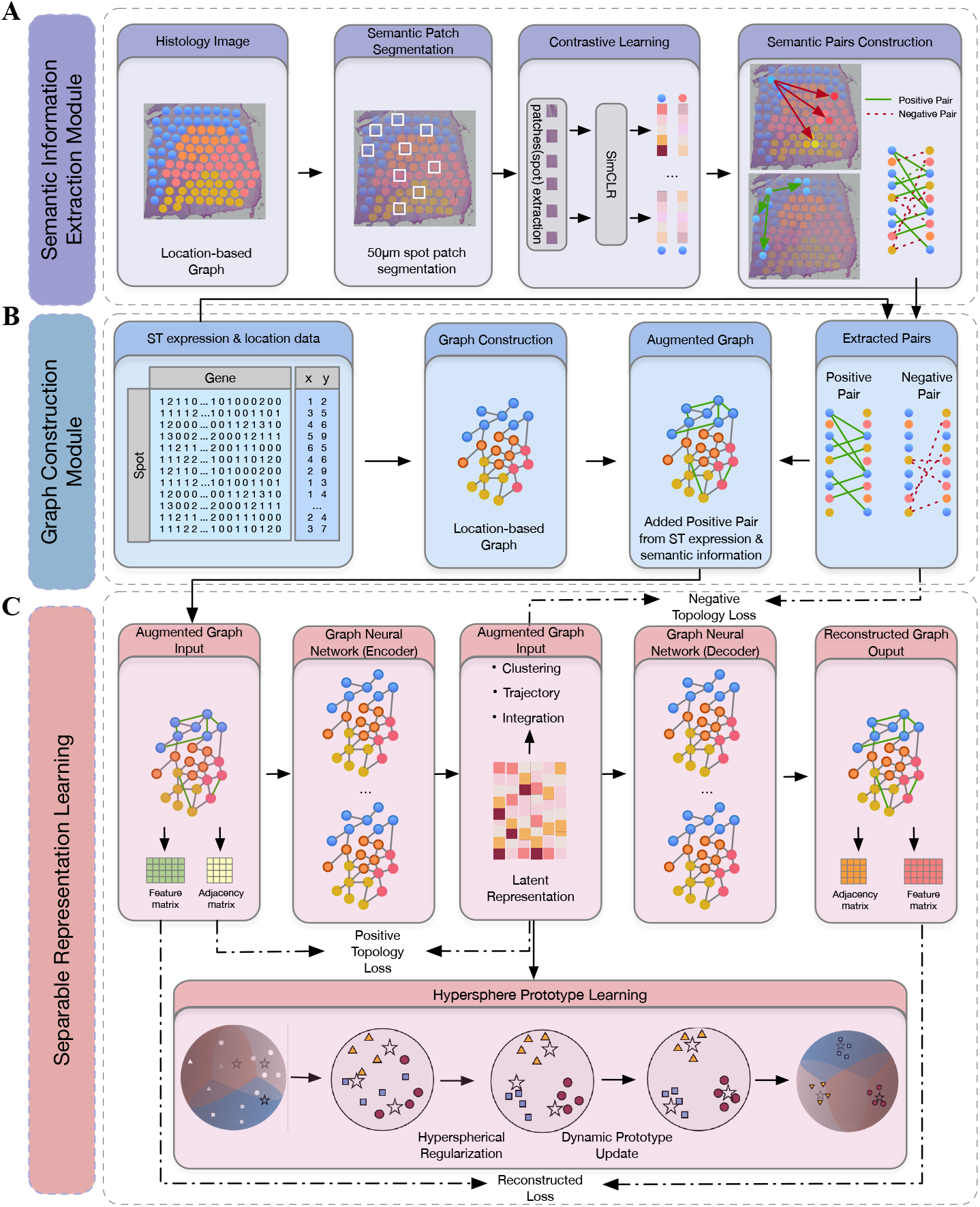
Overview of the S3RL framework for spatial transcriptomics analysis with three key modules. (A) **Semantic Information Extraction Module:** Histological images are segmented into spatially coherent regions based on a 50*µ*m spot segmentation approach. Then the contrastive learning-based SimCLR model is used to extract high-level semantic features from histological images, based on which positive/negative pairs with high/low semantic similarity are constructed. (B) **Graph Construction Module:** A location-based spatial *k*-nearest neighbor graph is first built from the ST data. Then this spatial graph is improved by incorporating the positive pairs with high semantic and gene expression similarity, enabling the capture of both local spatial structure and global semantic relationships between spots. (C) **Separable Representation Learning Module:** Graph neural networks (GNNs) are employed as the backbone (an encoder-to-decoder architecture) that collaboratively takes into account the spatial information (locations of spots), the semantic information (histological images of spots), and the gene expression profiles of spots in the spatial transcriptomics context. The encoder learns expressive low-dimension representations, which maintain the topological structure of extracted positive and negative pairs in the latent space, and the decoder is responsible for reconstructing the feature matrix. Subsequently, S3RL retrieves clustering results in an end-to-end manner via a hyperspherical prototype learning submodule. (D) **Hyperspherical Prototype Learning Submodule:** The representations are mapped and constrained to the unit-hypersphere with evenly distributed prototypes, where each prototype represents the center of a specific cell type. In this way, the affinity of each spot to the prototypes corresponding to different cell types is measured by the cosine similarity between the representations of the spot and prototypes. S3RL further introduces a hyperspherical regularization to ensure that all spots are evenly and closely distributed across prototypes and thus promote separability of the learned representations. Additionally, the current clustering results acting as pseudo labels provide a supervision signal to update the prototypes dynamically, which facilitates the adaptation of the prototypes to histological variations and thus improves the clustering performance.

**Fig. 2.**
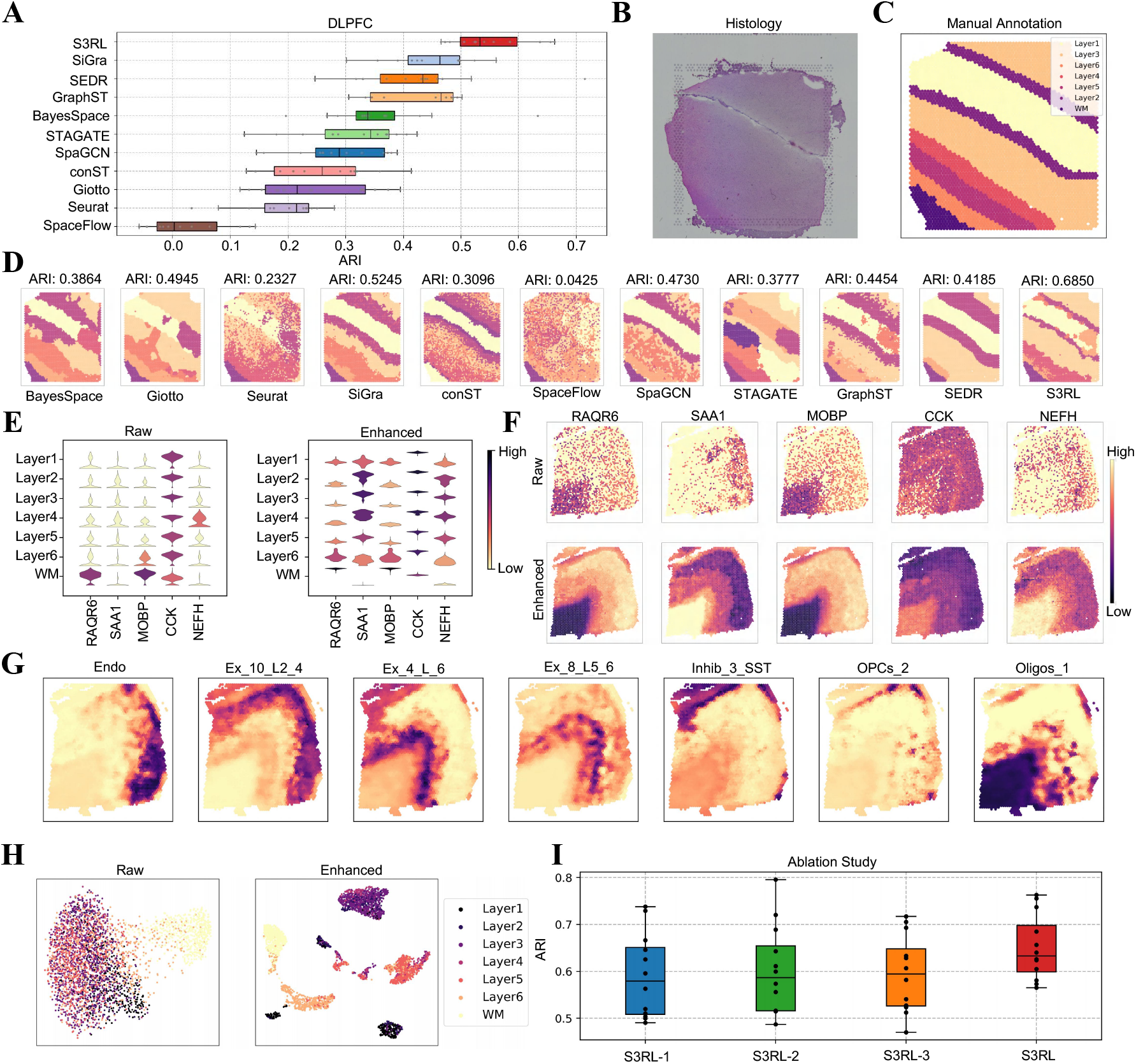
Spatial clustering and deconvolution analysis of human dorsolateral prefrontal cortex (DLPFC) using S3RL. (A) Comparison of Adjusted Rand Index (ARI) across different clustering methods on the DLPFC dataset. S3RL achieves the highest ARI of 0.57, outperforming other methods such as BayesSpace, Giotto, Seurat, SiGra, GraphST, and STAGATE. (B-C) Histology image (B) and manually annotated spatial layers (C) used as ground truth for clustering evaluation. (D) Clustering results from various methods on slice 151673, where S3RL achieves an ARI of 0.57, demonstrating improved spatial organization and boundary accuracy compared to alternative approaches. (E) Violin plots comparing raw and enhanced expression distributions for marker genes across cortical layers, showing improved inter-layer variability in enhanced data. I. Spatial expression maps of representative marker genes (e.g., PD4R6, SAA1, MOBP, CCK, NEFH), highlighting sharper gene expression boundaries in the enhanced data. (G) Deconvolution analysis using scRNA-seq and ST data on slice 151673, revealing spatial distributions of key cell types, including endothelial cells (Endo), excitatory neurons (Ex), inhibitory neurons (Inhib), oligodendrocyte precursor cells (OPCs), and oligodendrocytes (Oligos). (H) t-SNE visualization of clustering results, demonstrating improved spatial structure in S3RL-enhanced data. Ablation study results, showing ARI scores for different variants of S3RL, where the full model configuration achieves the highest clustering performance.

To further illustrate the performance of S3RL, we analyzed the clustering of slice 151673. S3RL achieves an ARI of 0.69, surpassing GraphST and STAGATE, while Giotto and Seurat achieved lower scores of 0.29 and 0.38, suffering from boundary overlaps.

We also examined gene expression patterns across cortical layers in both raw and S3RL-enhanced data (Figure 2E). The enhanced data provided a clearer distinction between layers, particularly where expression similarities exist. Additionally, Supplementary Fig.S3 compares the latent space clustering, where S3RL-enhanced embeddings exhibit greater compactness and improved layer separation.

To further assess spatial relationships among cortical layers, we conducted cell trajectory analysis across 12 slices of the 10X Visium DLPFC dataset using S3RL (Supplementary Fig. S4). Inferred inter-layer transitions were visualized as connectivity graphs, where each node represents a distinct anatomical layer (e.g., Layer1–Layer6 and WM), and edge thickness indicates the strength of interaction between layers. Notably, S3RL consistently identifies strong transitions between Layer4 and WM, as well as Layer1 and deeper cortical layers, across multiple slices. These consistent yet region-specific interaction patterns suggest that S3RL not only captures biologically meaningful spatial connectivity but also preserves inter-slice coherence in spatial architecture inference. This highlights the model’s ability to uncover spatial lineage relationships while maintaining robustness across heterogeneous tissue sections.

Furthermore, we visualized spatial gene expression of key markers, including PAQR6, SAA1, CCX, and MOBP [28] (Figure 2F) etc. S3RL-enhanced data better delineates expression boundaries compared to raw data(Figure 2H), aligning more accurately with known cortical structures.

To further evaluate the contributions of different modules within S3RL, we conducted an ablation study (Figure 2I). Specifically, we tested three ablated versions of the model: S3RL-1 does not use histology image-derived edges and does not update hyperspherical centers, S3RL-2 does not use histology image-derived edges but updates hyperspherical centers, and S3RL-3 incorporates image-derived edges but does not update hyperspherical centers. In contrast, the full S3RL model integrates both components. The results indicate that removing either component leads to a decrease in ARI scores, confirming that the complete configuration of S3RL achieves the best clustering performance. This suggests that both the image-derived edges and hyperspherical center updates are essential for optimizing spatial transcriptomics clustering.

### C. Deconvolution of scRNA-seq and ST Data in DLPFC Slice 151673

To further explore the integration of single-cell RNA sequencing (scRNA-seq) and spatial transcriptomics (ST) data, we conducted a deconvolution analysis on DLPFC slice 151673. We first applied S3RL to enhance the ST data, resulting in a spatial expression profile with clearer boundaries and improved expression fidelity. As the reference scRNA-seq data, we used a publicly available single-nucleus RNA-seq (snRNA-seq) dataset from the human dorsolateral prefrontal cortex (BA9) [29], generated by the 10x Genomics Chromium platform (GEO accession: GSE144136). This dataset comprises 78,886 nuclei and expression profiles of 30,062 genes.

As illustrated in Supplementary Fig. S5, the deconvolution framework draws inspiration from GraphST but adopts a simplified contrastive learning-based design. Specifically, we learn a projection matrix that maps scRNA-seq expression into the spatial domain. Unlike GraphST, which utilizes an autoencoder-reconstructed scRNA-seq matrix, our method directly uses the raw input matrix, which helps preserve intrinsic expression variability across cells and avoids potential information loss introduced by latent-space compression. The learned projection matrix transforms the scRNA-seq expression into a reconstructed spatial gene expression matrix, which is then aligned to the S3RL-enhanced ST matrix through contrastive learning. Positive and negative spot pairs are defined based on spatial proximity, enabling the model to maintain topological consistency while biologically estimating cell-type compositions at each spatial location. This framework allows biologically meaningful deconvolution of spatial transcriptomics guided by single-cell resolution information.

As shown in Figure 2G, our method effectively captures the spatial distributions of different cell types across the tissue, with distinct and well-defined regions corresponding to different cell types. The heatmaps illustrate the proportions of key cell types, including various excitatory neurons (Ex), oligodendrocytes (Oligos) and endothelial cells (Endo). Compared to the raw ST data, the deconvolved results provide a much clearer depiction of the underlying cellular architecture, highlighting the enriched presence of specific cell types in certain layers of the DLPFC. What is more, the results also achieves high spatial resolution and reveals biologically meaningful patterns. For example, the excitatory neurons (Ex) are predominantly found in the outer layers, while oligodendrocytes are concentrated in the white matter regions (Figure 2G) [30]. The deconvolution results demonstrate the ability of our method to accurately map the cell-type composition and provide a clearer representation of cellular heterogeneity in complex tissues like the DLPFC.

In addition, we present a qualitative comparison in Supplementary Fig. S6, where several representative reconstruction-based methods (e.g., STAGATE, SEDR, GraphST, and SiGra) are evaluated using the same deconvolution framework as ours. Although it is challenging to perform precise quantitative evaluation, visual inspection reveals that S3RL consistently produces the clearest expression boundaries and the most coherent spatial patterns. This further highlights the superior spatial consistency and structural fidelity of the representations learned by S3RL, offering strong support for its effectiveness in downstream deconvolution analysis.

### D. S3RL Improves Cell Type Clustering and Gene Expression Delineation on Nanostring Lung Slices

We also conducted a comprehensive analysis of the Nanostring dataset, which consists of 20 lung tissue slices [31]. Using the latent space representations learned by our model, we performed clustering and compared the results against several state-of-the-art spatial transcriptomics analysis methods (Figure 3A, Supplementary Fig. S7 presents the clustering results for all 20 slices in the latent space and Supplementary Fig. S8 presents the clustering results over all the methods across all the slices). The Adjusted Rand Index (ARI) was used as the evaluation metric. Our model achieved an average ARI of 0.6094 across 20 slices, significantly outperforming all other methods, with Seurat (the best-performing baseline) achieving 0.3989. This represents a relative improvement of approximately 52.8% over the best baseline and a 119.4% improvement over the average performance of all other methods. These results demonstrate our model’s superior ability to delineate cell types and tissue structures, providing a closer match to the ground truth labels (Figure 3B-D).

**Fig. 3.**
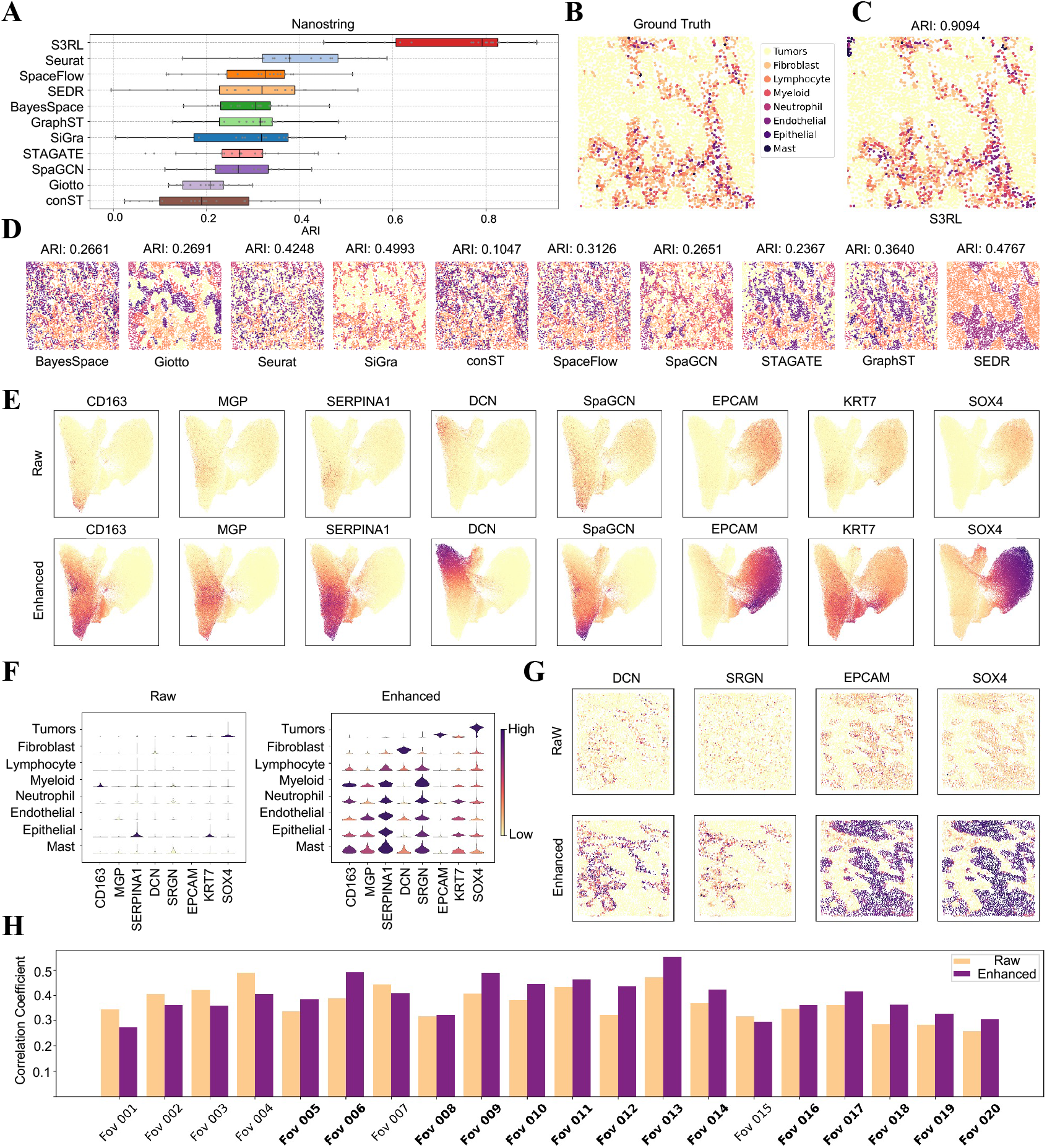
Nanostring Lung Slice Analysis and Correlation with TCGA Bulk RNA-seq. (A) Comparison of clustering performance across multiple methods using Adjusted Rand Index (ARI) on the Nanostring dataset. Our model achieves the highest ARI of 0.9094, significantly outperforming state-of-the-art methods such as Seurat, BayesSpace, SiGra, STAGATE, and GraphST. (B–C) Ground truth annotations (B) and the clustering result obtained using our model (C), demonstrating high spatial alignment with known cell types. (D) UMAP visualization of different clustering methods, showing clearer cluster separation in our enhanced data representation. (E) Spatial gene expression plots for selected markers (CD163, EPCAM, SOX4, etc.) before and after enhancement. The enhanced data exhibits sharper expression boundaries, revealing improved biological signal resolution. (F) Violin plots comparing raw and enhanced gene expression across different cell types, highlighting increased expression specificity in the enhanced data. (G) Spatial distribution of selected marker genes (EPCAM, DCN, SRGN, SOX4) across individual lung slices, where enhancement improves tissue-specific gene localization. (H) Correlation analysis between TCGA bulk RNA-seq data and Nanostring spatial transcriptomics (ST) data across 20 slices. Each bar represents the Pearson correlation coefficient between the bulk data and either raw (orange) or S3RL-enhanced (purple) spatial data. Slices where the enhanced data outperforms the raw are highlighted in bold, indicating improved alignment with bulk gene expression profiles in 14 out of 20 slices.

To further evaluate the model’s performance, we conducted a comparison of raw and enhanced data on the Nanostring dataset using slice 014. Our model achieved the highest ARI of 0.9094, significantly outperforming other methods (Figure 3D). The UMAP visualization plot of gene expression across all 20 slices (Figure 3E) shows that the S3RL-enhanced data exhibits a clearer separation of clusters compared to raw data, indicating improved resolution of cell-type heterogeneity (Supplementary Fig. S8 provides the clustering results across all 20 slices of all the compared methods). In addition, these plots (Figure 3E) also demonstrate that key marker genes such as CD163, EPCAM, and SOX4 exhibit sharper boundaries in the enhanced data, highlighting S3RL’s ability to amplify biologically relevant signals. Violin plots (Figure 3F) further emphasize these improvements, showing significant differences in gene expression patterns between raw and enhanced data.

For a more detailed examination, we selected one individual slice (slice 14) and analyzed the spatial distribution of genes EPCAM, DCN, SRGN, and SOX4. As shown in Figure 3G, enhanced data provides clearer spatial patterns, confirming our model’s improved sensitivity and specificity in capturing critical biological signals within lung tissue samples. Notably, EPCAM is highly expressed in lung tumors and serves as a biomarker for epithelial tumor cells [32]. In contrast, DCN acts as a tumor suppressor, with reduced expression correlating with poor prognosis in lung adenocarcinoma [33]. Additionally, SOX4 has been implicated in promoting epithelial-to-mesenchymal transition and tumor progression in lung cancer [34]. In addition, Supplementary Fig. S9 compares the gene expression reconstructions of these marker genes produced by several representative generative methods, including GraphST, SEDR, STAGATE, and SiGra. These methods each reconstructed the data based on their own model outputs. From the visualization results, S3RL shows overall clearer spatial localization and sharper expression boundaries, indicating stronger reconstruction fidelity. While STAGATE produces comparable reconstructions for most genes, S3RL exhibits a more pronounced contrast in the spatial pattern of *SRGN*, demonstrating improved distinguishability in marker gene recovery. What’s more, Supplementary Fig. S10 displays the UMAP visualizations of latent embeddings across all 20 Nanostring lung slices, comparing raw and S3RL-enhanced representations. The enhanced data reveals more compact and well-separated clusters, indicating improved spatial structure and cell-type distinguishability within the latent space.

Furthermore, Supplementary Fig. S11 presents the spatial distribution of these four marker genes (EPCAM, DCN, SRGN, and SOX4) across all 20 Nanostring slices, comparing both raw and S3RL-enhanced data. The enhancement results consistently demonstrate improved gene expression resolution, reinforcing the effectiveness and generalizability of S3RL across multiple tissue samples.

### E. Correlation Analysis Between TCGA Lung Cancer Bulk RNA-seq and Nanostring ST Data

We then evaluated the correlation between TCGA bulk RNA-seq data (TCGA-LUAD) and single-cell spatial transcriptomics (ST) data from the TCGA database and Nanostring dataset to assess our model’s effectiveness in capturing biologically relevant expression patterns. Pearson correlation coefficients were calculated for each slice, comparing raw and enhanced ST data against bulk RNA-seq profiles. As shown in Figure 3H, the enhanced data yields higher correlation values in 14 out of 20 slices(over 70%), demonstrating that S3RL more effectively aligns spatial transcriptomic signals with bulk tissue-level expression patterns.

To further validate this finding, we compared our method with four representative spatial transcriptomics methods that include data reconstruction capabilities—GraphST, SEDR, SiGra, and STAGATE. Supplementary Fig. S12 presents the correlation bar plots for each method across all 20 slices, comparing their reconstructed outputs (enhanced) and raw ST data. Notably, our S3RL-enhanced data consistently achieves higher correlation values than the corresponding raw data in nearly all slices. In contrast, SEDR and SiGra show little to no improvement, while STAGATE exhibits modest gains.

Interestingly, GraphST displays a unique pattern: its reconstructed data yields higher correlation values than raw data across all slices. However, as shown in Supplementary Fig. S9, the reconstructed spatial expression of marker genes by GraphST appears overly smoothed and lacks distinct expression boundaries. We hypothesize that the inflated correlation values may result from this over-smoothing effect, where uniform expression values lead to artificially high similarity with bulk profiles despite reduced biological interpretability. In contrast, S3RL maintains sharper gene-specific expression patterns while simultaneously improving alignment with bulk RNA-seq data, underscoring its strength in balancing biological resolution and statistical correlation.

To further investigate this trend, we analyzed the scatter plots of correlation values between bulk RNA-seq and spatial transcriptomics for each field of view (FOV), as shown in Supplementary Fig. S13 and Fig. S14. While the raw ST data exhibits a more concentrated distribution along the diagonal, its overall correlation with bulk RNA-seq is lower. This suggests that although some genes in the raw data match bulk RNA-seq values more closely, the overall expression patterns remain suboptimal due to noise and data sparsity. In contrast, the enhanced ST data demonstrates a higher correlation coefficient, signifying a more consistent alignment with bulk RNA-seq expression trends. However, the scatter plots reveal that enhanced data points exhibit slightly increased dispersion, indicating that while the global consistency with bulk RNA-seq improves, local variability remains, likely due to the non-linear transformation applied during enhancement.

These findings further validate that our model enhances the consistency between bulk RNA-seq and single-cell ST data, thereby bridging the gap between bulk and single-cell perspectives. The enhancement process improves global gene expression alignment while maintaining meaningful biological variability, ultimately providing a more comprehensive and biologically interpretable representation of the tumor microenvironment in lung cancer tissues.

### F. Differential Gene Expression and Cell-Cell Communication in DLPFC Slice 151509

To further explore whether S3RL’s data representation and model reconstruction could more accurately capture and recover valuable signals and hidden biological meanings that are often overlooked due to sparse and noisy single-cell data, we chose to observe two distinct datasets in the following two subsections. Specifically, we selected slices from two datasets and analyzed their gene expression and cell-cell communication patterns both in the raw data and in the S3RL-enhanced reconstructed data. In the case of the DLPFC 151509 slice, we focused on differential gene expression across cortical layers and evaluated the effectiveness of S3RL-based data enhancement in improving spatial transcriptomic resolution.

As shown in Figure 4A, marker genes were identified across different cortical layers, and volcano plots were used to compare their expression between raw data and S3RL-reconstructed data. The enhanced data exhibited a stronger layer-specific expression pattern for key genes such as FGFR1, FGFR2, NOTCH4, and DLL4. For instance, FGFR1/FGFR2 were significantly expressed in Layer 1, Layer 5, and Layer 6, whereas DLL4/NOTCH4 showed a pronounced preference for Layer 3 and Layer 5. These results demonstrate that S3RL effectively refines biologically meaningful gene expression patterns, offering an improved representation of spatially resolved transcriptional activity.

**Fig. 4.**
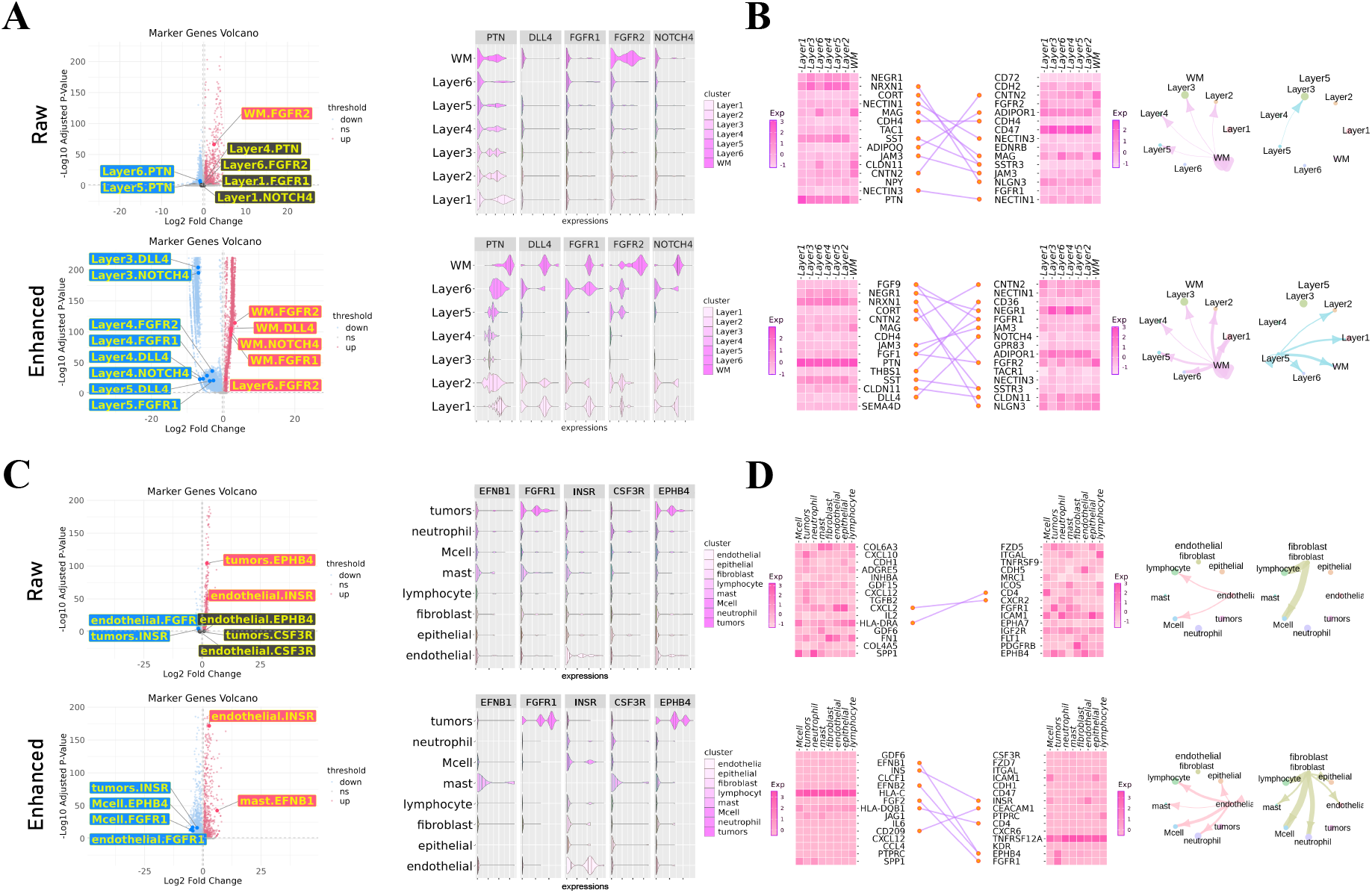
Differential Gene Expression and Cell-Cell Communication in DLPFC Slice 151509 and Nanostring Lung Cancer Slice 001. (A) Volcano plots and violin plots for differential gene expression analysis in DLPFC slice 151509, comparing raw and S3RL-enhanced data. The enhanced data reveals more layer-specific expression of key marker genes such as FGFR1, FGFR2, DLL4, and NOTCH4, along with clearer stratification across cortical layers. (B) Cell-cell communication analysis of the same DLPFC slice. The heatmaps, interaction plots, and chord diagrams show enhanced, particularly signaling in the FGF and Notch pathways after applying S3RL. (C) Volcano plots and violin plots for Nanostring lung cancer slice 001, showing improved localization and signal clarity for genes like INSR, EPHB4, and FGFR1 in the enhanced data, particularly within tumor and immune-related clusters. (D) Cell-cell communication analysis of the same lung slice. The heatmaps and interaction networks illustrate more pronounced tumor-immune signaling, highlighting enhanced detection of the INS–INSR, FGF–FGFR, and EFNB–EPHB pathways following S3RL enhancement.

To further investigate the impact of S3RL enhancement, violin plots were generated to visualize expression distributions of these key genes before and after reconstruction. In raw data, some genes exhibited diffuse or ambiguous distribution across layers, making it difficult to delineate layer-specific patterns. However, in S3RL-enhanced data, genes such as FGFR1, FGFR2, and NOTCH4 displayed sharper layer distinctions, demonstrating S3RL’s ability to enhance spatial resolution. We further employed CellChat [12] to infer ligand-receptor interactions between cortical layers. The heatmap illustrates the intensity of ligand-receptor interactions, showing increased expression levels for key receptor-ligand pairs such as DLL4-NOTCH4 and FGF-FGFR axes in the enhanced data (Figure 4B). Additionally, network diagrams highlight inter-layer communication patterns, where S3RL-enhanced data reveal more pronounced inter-layer signaling, particularly between Layer 3, Layer 5, and white matter (WM), further validating S3RL’s capacity to reconstruct intercellular communication networks with greater accuracy.

Building upon these findings, we examined two major signaling pathways in cortical organization. FGF8/FGF16-FGFR1/FGFR2 (Fibroblast Growth Factor Pathway) is critical for GnRH neuron maturation and plays a key role in tumor cell survival [35]–[37]. In S3RL-enhanced data, FGFR1 and FGFR2 showed stronger expression in Layer 1, Layer 5, and Layer 6, indicating S3RL’s ability to refine FGF signaling representation in cortical layers. Similarly, DLL4-NOTCH4 signaling is essential for cerebral vascularization and blood-brain barrier maintenance, and it plays a crucial role in glioblastoma angiogenesis [38], [39]. In S3RL-enhanced data, DLL4 and NOTCH4 exhibit clearer expression in Layer 3 and Layer 5, confirming S3RL’s improved reconstruction of Notch-mediated vascular formation. These results collectively demonstrate S3RL’s effectiveness in refining cortical layer-specific gene expression and cell-cell communication networks, providing a powerful framework for enhancing spatial transcriptomics in complex brain tissues.

### G. Differential Expression and Cell-Cell Communication in Nanostring Lung Cancer Slice 003

To investigate tumor-specific transcriptional patterns and intercellular signaling in lung tissue, we further applied S3RL enhancement to the Nanostring slice 003. This allowed for a more precise identification of differentially expressed genes across spatial clusters and a clearer characterization of cell-cell communication events within the tumor microenvironment. As shown in Figure 4C, volcano plots compare gene expression patterns between raw and S3RL-reconstructed data. The enhanced data exhibits stronger cluster-specific expression for key genes, such as INSR, FGFR1, EPHB4, and CSF3R, with INSR enriched in endothelial cells, FGFR1 in macrophages (Mcell), and EPHB4 in tumor cells. These findings suggest that S3RL effectively sharpens the spatial resolution of key gene expressions, enhancing the biological interpretability of the data.

To further explore gene expression distribution, violin plots were generated to illustrate the spatial localization of FGFR1, INSR, CSF3R, and EPHB4 across different clusters. Compared to raw data, the S3RL-enhanced dataset reveals clearer expression patterns, including increased FGFR1 expression in macrophages, INSR enrichment in tumor and endothelial cells, and more pronounced EPHB4 expression in tumor cells. Additionally, we employed CellChat [12] to construct cell-cell communication networks and used heatmaps to quantify ligand-receptor interactions between clusters (Figure 4D). The enhanced data reveals a significant increase in biologically meaningful intercellular interactions, particularly in the INS-INSR, FGF-FGFR, and EFNB-EPHB signaling pathways. Network diagrams further illustrate strengthened communication between tumor cells, endothelial cells, and immune cells (mast, lymphocyte, and neutrophil). Notably, fibroblasts also show increased interactions with neutrophils, mast cells, and lymphocytes, suggesting an enhanced immune regulatory role within the tumor microenvironment.

Building upon these findings, we further investigated the oncogenic mechanisms of the INS-INSR, FGF2-FGFR1, and EFNB1/EFNB3-EPHB4 signaling pathways. The INS-INSR axis is overexpressed in endothelial cells and serves as a hallmark of tumor vascularization [40]. The S3RL-enhanced data confirms a significant upregulation of INSR in tumor and endothelial cells, highlighting its role in tumor angiogenesis and tumor-endothelial interactions. The FGF2-FGFR1 axis, a key fibroblast growth factor signaling pathway, is known to drive cancer cell proliferation, angiogenesis, and metastasis [36], [41]. In the enhanced dataset, FGFR1 expression is markedly increased in macrophages and tumor cells, consistent with FGF-FGFR signaling aberrations observed in NSCLC, breast cancer, and gastrointestinal cancers [42]. Furthermore, the EFNB1/EFNB3-EPHB4 axis, a receptor tyrosine kinase (RTK) pathway, plays a crucial role in promoting tumor survival, migration, and vascularization [43], [44].In the S3RL-enhanced data, EPHB4 expression is significantly upregulated in tumor cells (violin plot in Fig.4B), consistent with a 3-fold increase in lung cancer tissues compared to normal tissues [45], and our data show enhanced interaction with immune cells, particularly mast and lymphocyte clusters. These results demonstrate that the S3RL method improves the resolution of key oncogenic signaling pathways, enhancing the biological relevance of spatial transcriptomics in tumor microenvironment studies.

### H. Comparison Between S3RL and SiGra in L-R Pair Identification and Cell-Cell Communication Analysis

Accurate identification of ligand-receptor (L-R) interactions is essential for deciphering cell-cell communication in single-cell spatial transcriptomics. To construct a comprehensive and high-confidence L-R interaction set, we integrated candidate L-R pairs from SiGra [20], which aggregates interactions from multiple databases, and the updated International Union of Pharmacology (IUPHAR) database [46]. After removing redundancies and ensuring uniqueness, we curated a final set of 2864 reliable L-R interaction pairs, providing a robust foundation for downstream analyses of intercellular signaling dynamics. Among them, genes of 480 L-R pairs are included in this NanoString CosMx dataset (189 ligands and 159 receptors).

In Supplementary Fig. S15, we compared S3RL against four representative enhancement-capable methods (STAGATE, GraphST, SEDR, and SiGra) in L-R pair significance analysis. To further quantify their capabilities in detecting functionally meaningful L-R interactions, we further introduced three evaluation metrics and visualized them using a radar plot (bottom panel of Supplementary Fig. S15): the Enhanced Specificity Ratio, Effective Enhancement Percentage, and Specificity Stability Percentage. These metrics assess the tradeoff between discovering new specific interactions and preserving biologically important existing ones. Results show that both S3RL and STAGATE achieve the strongest overall performance, which further confirm the robustness of S3RL in resolving cell-cell communication within spatial transcriptomic data.

Take SiGra method as an example, our proposed S3RL approach enhances data quality by identifying a greater number of biologically meaningful L-R pairs while effectively reducing false positives (Supplementary Fig. S15). In the L-R pair significance analysis (scatter plot), S3RL significantly increases the number of shared L-R pairs (orange, 452 vs. 446 in SiGra) and specifically enhanced L-R pairs (green, 21 vs. 16 in SiGra), demonstrating its capacity to reveal previously undetected but biologically relevant communication patterns. Furthermore, S3RL identifies two lung cancer-associated L-R interactions, ANXA1–FPR1 and OSM–LIFR, which are absent in SiGra’s results. The ANXA1–FPR1 pathway is implicated in tumor immune evasion via immuno-suppressive microenvironment formation [47], while OSM–LIFR signaling contributes to tumor progression through STAT3 activation and EMT induction [48]. Additionally, S3RL detects a large number of L-R pairs at the upper right corner (−log_10_ FDR ≈ 10), indicating that these ligand-receptor interactions exhibit extremely high statistical significance (FDR close to zero) in the enhanced data, further validating their biological relevance. Moreover, other competitors retains a greater number of false-positive L-R pairs in the lower right quadrant, where interactions are significant in the raw data but become non-significant after enhancement. This suggests that these methods may still preserve low-confidence signals, whereas S3RL more effectively eliminates such noise, leading to a more precise L-R interaction identification.

### I. S3RL Accurately Segments Neural and Tumor Tissues and Enhances Biological Interpretation in Mouse Brain and Human Breast Cancer

We further evaluated the applicability of S3RL on 10X Visium spatial transcriptomics datasets, including Mouse Brain Anterior and Human Breast Cancer. As shown in Figure 5, our results demonstrate that S3RL exhibits high accuracy in spatial tissue segmentation across distinct biological samples, closely aligning with ground truth annotations. In the Mouse Brain Anterior dataset (Figure 5A), S3RL achieved an Adjusted Rand Index (ARI) of 0.51, effectively capturing the spatial organization of distinct brain regions, including cortical and subcortical zones, validating its effectiveness in neural tissue analysis. Similarly, in the Human Breast Cancer dataset (Figure 5B), S3RL obtained an ARI of 0.67, demonstrating strong performance in delineating tumor microenvironments, successfully segmenting tumor, stromal, and other critical structural regions, providing deeper insights into breast cancer tissue architecture. To further evaluate the performance of S3RL relative to other approaches, we compared its clustering accuracy with several representative methods that possess spatial reconstruction capabilities, including GraphST, SiGra, STAGATE, and SEDR. As illustrated in Figure 5C, S3RL consistently outperforms or matches these methods across both datasets (detailed clustering results are shown in Supplementary Fig. S16). The radar plot summarizes the ARI scores, normalized to the performance of S3RL, and highlights its robust generalization and effectiveness in accurately segmenting spatial domains across diverse tissue types.

**Fig. 5.**
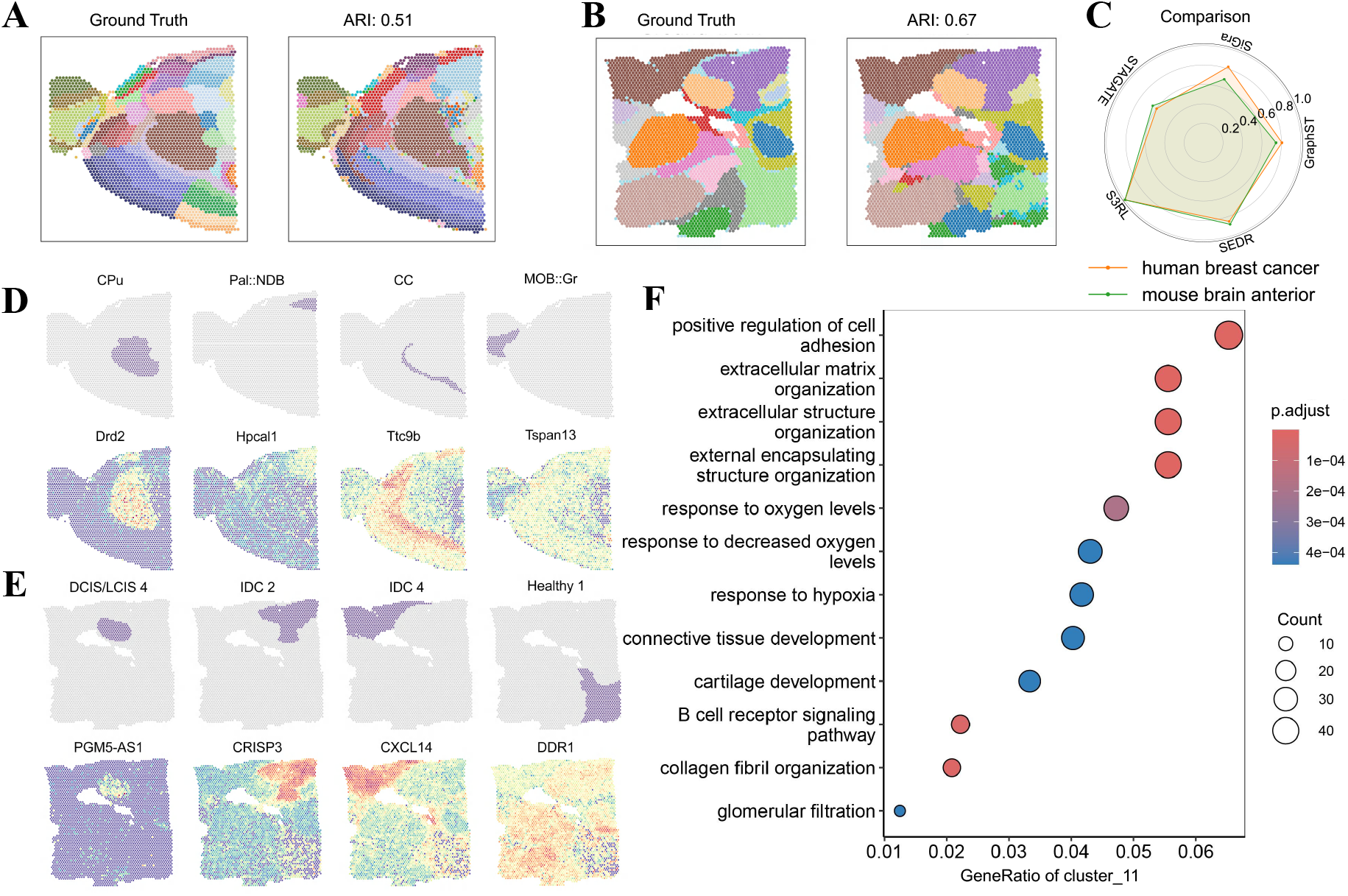
Evaluation of S3RL on Mouse Brain Anterior and Human Breast Cancer Datasets. (A) Segmentation results on the Mouse Brain Anterior dataset, comparing ground truth annotations and S3RL-predicted spatial domains, achieving an Adjusted Rand Index (ARI) of 0.51. The model effectively captures cortical and subcortical structures, confirming its accuracy in neural tissue segmentation. (B) Segmentation performance on the Human Breast Cancer dataset, where S3RL achieved an ARI of 0.67, accurately distinguishing tumor, stromal, and other structural regions within the microenvironment. (C) Radar plot comparison of ARI scores across five methods (GraphST, SiGra, STAGATE, S3RL, and SEDR) on both datasets. Each method’s performance is normalized against S3RL (serving as the reference with a value of 1.0). (D) Spatial expression of key marker genes (Drd2, Hpcal1, Ttc9b, Tspan13) in the Mouse Brain Anterior dataset, illustrating region-specific gene localization. (E) Spatial gene expression patterns in the Human Breast Cancer dataset for PGM5-AS1, CRISP3, CXCL14 and DDR1, highlighting distinct tissue-specific gene enrichment. (F) Pathway enrichment analysis in the Human Breast Cancer dataset, showing significantly enriched pathways such as “extracellular matrix organization,” “response to hypoxia,” and “B cell receptor signaling pathway.” The dot plot visualizes gene ratios across clusters, emphasizing S3RL’s ability to enhance biologically relevant signal detection. These results collectively validate S3RL’s efficacy in segmenting tissue architecture and refining functional insights in spatial transcriptomics.

To further validate S3RL’s ability to enhance biological feature resolution, we visualized the spatial expression of key marker genes. As shown in Figure 5D and Figure 5E, in the Mouse Brain dataset, genes such as Drd2, Hpcal1, Ttc9b, and Tspan13 exhibit clear spatial distribution across different brain regions. Notably, Drd2 is highly expressed in the caudoputamen (CPu), consistent with its key role in dopaminergic signaling in the striatum [49]. Hpcal1, on the other hand, is a neuronal calcium-binding protein belonging to the visinin-like protein (VILIPs) family, which is broadly expressed in the nervous system and plays a role in calcium-mediated neuronal signaling. Notably, Pal::NDB, a region of the basal forebrain, is functionally connected to the hippocampus via cholinergic projections, and given the involvement of calcium-binding proteins in synaptic modulation, Hpcal1 may have a regulatory role in this cross-regional interaction. In the Human Breast Cancer dataset, genes like PGM5-AS1, CRISP3, CXCL14, and DDR1 show enhanced enrichment patterns across tissue structures. Notably, CXCL14 is known for its tumor-suppressive properties in breast cancer, with studies indicating its ability to inhibit tumor growth and metastasis [50]. Similarly, DDR1 overexpression has been linked to tumor progression and poor prognosis in breast cancer patients [51]. These consistent expression patterns across different tissue regions highlight their potential as reliable biomarkers for breast cancer characterization. Compared to raw data, the enhanced spatial maps generated by S3RL provide clearer spatial gene expression signatures, demonstrating the model’s ability to amplify biologically meaningful signals.

Additionally, pathway enrichment analysis in the Human Breast Cancer dataset (Figure 5F) identified key pathways associated with tumor microenvironment and cancer progression, including “extracellular matrix organization,” “response to hypoxia,” and “B cell receptor signaling pathway.” The dot plot reveals significant enrichment for these pathways, with high gene ratios in specific regions, highlighting the biological relevance of S3RL-enhanced data. These results collectively show that S3RL not only improves spatial tissue segmentation across diverse biological datasets but also enhances gene expression signals, refines cell type annotations, and provides deeper biological insights, reinforcing its robustness in spatial transcriptomics analysis.

### J. S3RL Reveals Region-Specific Functional Domains in Mouse Olfactory Bulb and Hippocampus through Spatial Gene Annotation

We applied S3RL to segment and annotate spatial domains in the mouse olfactory bulb (Stereo-seq data) and mouse hippocampus (Slide-seqV2 data)(Figure 6), utilizing DAPI-stained images for reference. Marker gene expression (Figure 6C and Figure 6F) and GO enrichment analysis (Figure 6G and Figure 6H) validated the identified spatial regions, highlighting biologically relevant processes in each dataset.

**Fig. 6.**
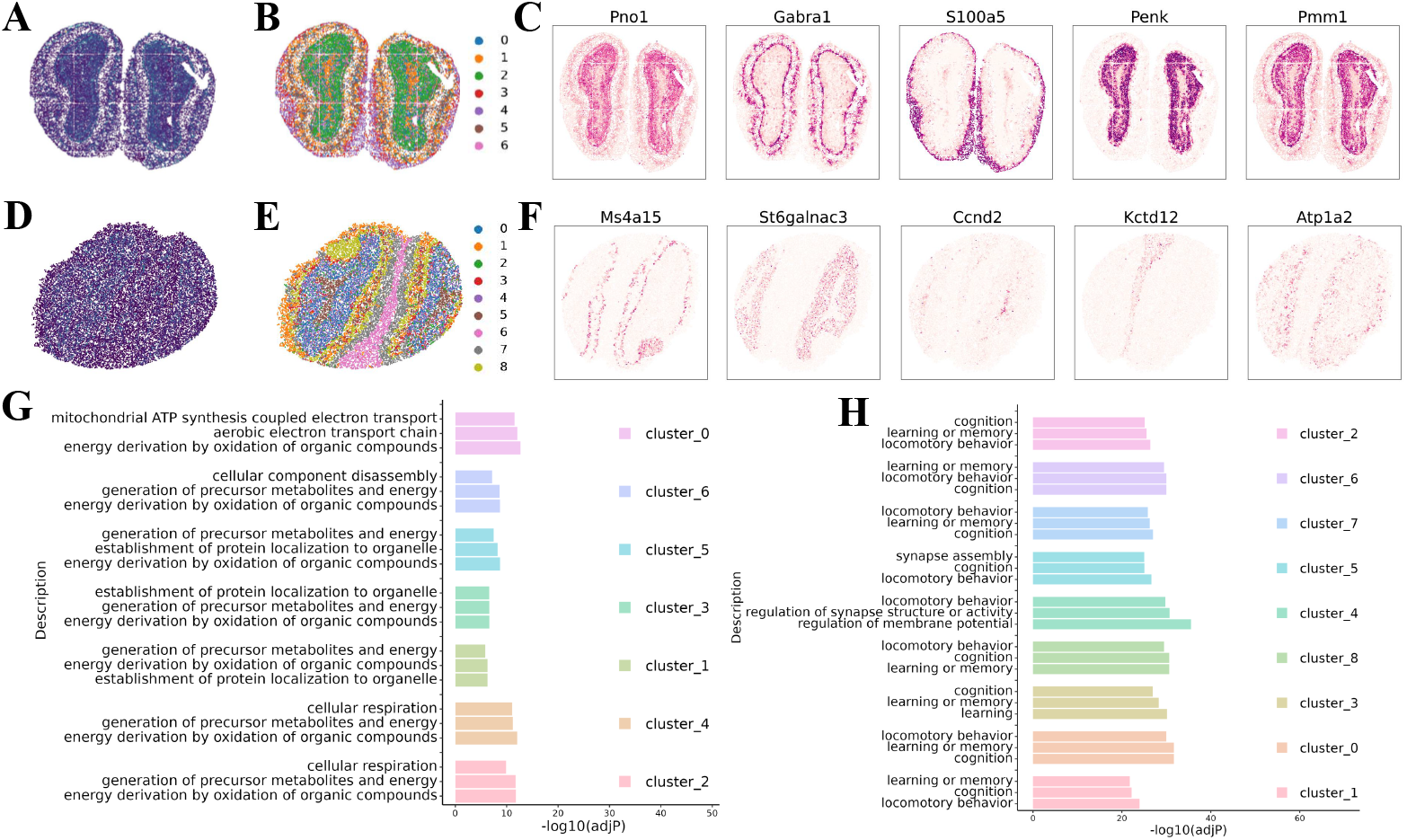
Spatial Analysis of Mouse Olfactory Bulb and Hippocampus using S3RL. (A) Raw spatial transcriptomics data of the mouse olfactory bulb, showing spatially organized cell populations. (B) Spatial domain segmentation obtained using S3RL, identifying distinct functional regions within the olfactory bulb. (C) Expression patterns of selected marker genes (Pno1, Gabra1, S100a5, Penk, and Pmm1) highlighting the structural organization of the olfactory bulb. (D) Raw spatial transcriptomics data of the mouse hippocampus, illustrating the spatial arrangement of cell populations. (E) Spatial domain segmentation in the hippocampus using S3RL, capturing biologically meaningful subregions. (F) Expression patterns of key marker genes (Ms4a15, St6galnac3, Ccnd2, Kctd12, and Atp1a2) within distinct hippocampal subregions. (G) Gene ontology (GO) enrichment analysis for identified spatial clusters in the olfactory bulb, showing significant enrichment in mitochondrial ATP synthesis, aerobic electron transport chain, and precursor metabolite generation (cluster 0, cluster 6, and cluster 5), reflecting the metabolic demands of olfactory sensory neurons. Additional enrichments in cellular component disassembly and organelle localization (cluster 3 and cluster 1) suggest synaptic remodeling and neuronal turnover. (H) GO enrichment analysis for hippocampal clusters, revealing significant enrichment in cognition, learning and memory, and synapse assembly (cluster 2, cluster 6, and cluster 7), consistent with hippocampal neuroplasticity. Locomotory behavior and regulation of synapse structure (cluster 4 and cluster 8) further support the hippocampus’s role in spatial navigation. These results demonstrate S3RL’s ability to accurately resolve spatially distinct biological processes in complex neural tissues.

We applied S3RL to segment and annotate spatial domains in the mouse olfactory bulb (Stereo-seq data) and mouse hippocampus (Slide-seqV2 data) (Figure 6), utilizing DAPI-stained images for reference (Figure 6A and D). The segmentation results (Figure 6B and E) demonstrate distinct spatial domains, which were further validated through marker gene expression (Figure 6C and F) and GO enrichment analysis (Figure 6G and H), highlighting biologically relevant processes in each dataset. In the olfactory bulb, marker genes such as Pno1, Gabra1, S100a5, Penk, and Pmm1 (Figure 6C) delineate the structural organization, with Gabra1 encoding a GABA-A receptor subunit critical for inhibitory neurotransmission in the olfactory bulb [52], and S100a5 and Penk associated with neuronal differentiation and signaling regulation, respectively [53], [54]. These genes collectively highlight the layered organization of the olfactory bulb, including the glomerular and mitral cell layers, consistent with its role in olfactory processing. In the hippocampus, expression patterns of key marker genes such as Ms4a15, St6galnac3, Ccnd2, Kctd12, and Atp1a2 (Figure 6F) distinguish subregions such as the dentate gyrus (DG), CA1, CA2, and CA3. For instance, Ccnd2, involved in cell cycle regulation, is enriched in the DG, supporting its role in adult neurogenesis [55], while Atp1a2, encoding a Na+/K+ ATPase subunit, maintains neuronal ion gradients across hippocampal subregions [56]. These marker genes underscore the functional specialization of hippocampal subregions, with implications for learning and memory processes. These findings demonstrate that S3RL effectively captures region-specific gene expression patterns, providing insights into the functional organization of the mouse olfactory bulb and hippocampus.

Leveraging S3RL for spatial clustering, we conducted GO enrichment analysis to characterize the biological functions of the identified regions. In the mouse olfactory bulb, GO analysis revealed enrichment in mitochondrial ATP synthesis, aerobic electron transport chain, and precursor metabolite generation, particularly in cluster 0, cluster 6, and cluster 5 (Figure 6G)). These findings align with the high metabolic demands of olfactory sensory neurons, essential for continuous odor processing [57]. Additionally, cellular component disassembly and organelle localization were enriched in cluster 3 and cluster 1, reflecting neuronal turnover and synaptic remodeling in response to sensory stimuli.

For the mouse hippocampus, GO analysis (Figure 6H)) identified cognition, learning and memory, and synapse assembly as significantly enriched in clusters such as cluster 2, cluster 6, and cluster 7, consistent with the hippocampus’s role in neuroplasticity. Regulation of synapse structure and locomotory behavior in cluster 4 and cluster 8 further supports its involvement in spatial navigation. These results also demonstrate S3RL’s capability to resolve spatially distinct biological processes, enabling precise functional mapping in complex neural tissues.

### K. Clustering Performance under Multi-Slice Alignment

In spatial transcriptomics analysis, aligning and integrating multiple tissue slices is critical for ensuring data consistency and improving the interpretability of spatial patterns. Among existing methods, GraphST is one of the few that supports multi-slice alignment. Therefore, we selected GraphST as the baseline method for comparison in this setting.

However, GraphST is limited to aligning only fully paired slices, while our proposed method, S3RL, is compatible with both complete and partially missing tissue slices. To comprehensively evaluate clustering robustness, we assess the performance of S3RL and GraphST under two alignment scenarios: full-slice alignment and partial-slice alignment. This comparison highlights the flexibility of S3RL in handling incomplete spatial transcriptomics data, which is common in real-world biological samples.

In our approach, aligned tissue slices are embedded into a unified coordinate system, enabling joint clustering across slices. This spatial unification, facilitated by our graph construction design, ensures that spatially similar spots from different slices are effectively aligned and incorporated into the same spatial graph. Consequently, the local neighborhood of each spot can benefit from additional context provided by aligned adjacent slices, further improving the accuracy and consistency of downstream clustering results.

#### 1) Clustering Performance on Complete Slices

We first selected four complete slices from the 10X Visium dataset and performed alignment at the initial stage of data processing. The aligned slices were then integrated to enhance the utilization of neighboring cells before representation learning and clustering. As shown in Figure 7 (A, B, C), the alignment performance of S3RL is superior to that of GraphST, ensuring that similar cells from different slices are more accurately matched. In contrast, GraphST exhibits weaker alignment in certain regions, leading to less effective data integration.

**Fig. 7.**
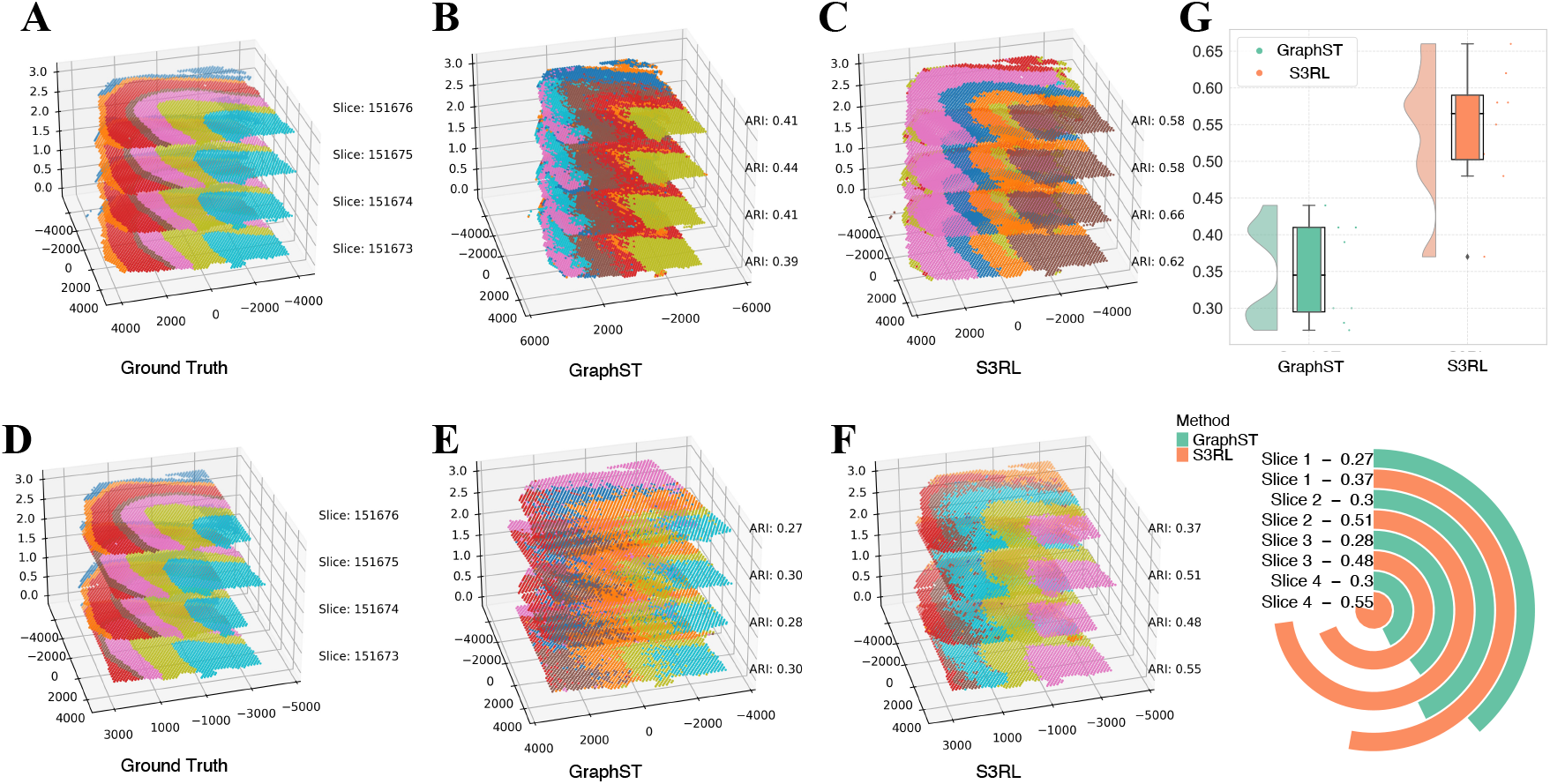
Multi-Slice Clustering Performance Evaluation. (A) Ground truth spatial domains of multiple tissue slices, illustrating the expected spatial organization. (B) Clustering results obtained using GraphST, where Adjusted Rand Index (ARI) values indicate the alignment accuracy across slices. GraphST exhibits lower ARI scores and reduced spatial consistency. (C) Clustering results obtained using S3RL, demonstrating higher ARI scores and improved spatial consistency across slices. The S3RL model effectively preserves spatial continuity and enhances cluster separability. (D) Ground truth spatial domains for another set of slices, serving as a reference for evaluating clustering performance. (E) GraphST clustering results on the second dataset, showing lower ARI values and weaker spatial alignment. (F) S3RL clustering results on the second dataset, with higher ARI values and superior spatial structure preservation. (G) Performance comparison between GraphST and S3RL. The raincloud plot (top right) illustrates the ARI distribution across slices, where S3RL consistently achieves higher clustering accuracy. The circular bar chart (bottom right) further quantifies slice-wise ARI scores, highlighting S3RL’s superior performance across all evaluated slices. These results confirm that S3RL effectively integrates multi-slice spatial transcriptomics data, ensuring robust clustering and improved tissue structure preservation.

Furthermore, the raincloud plots Figure 7G illustrate the clustering performance of different methods. These plots, which combine violin and box plots, provide insights into the distribution characteristics of the clustering results. S3RL demonstrates more compact and well-separated clusters, indicating lower intra-cluster variance and better-defined cluster boundaries. In contrast, GraphST exhibits higher dispersion in the clustering results, suggesting less precise cluster assignments.

#### 2) Robustness to Partial Slice Removal

To simulate real-world scenarios where slices may be incomplete or have limited overlap, we randomly cropped 30% of each slice and applied the same alignment, representation learning, and clustering process. This setup introduces greater complexity and challenges, making it crucial to assess the robustness of our method.

As depicted in Figure 7 (D, E, F), S3RL maintains a high level of alignment accuracy even when 30% of the slices are removed. In contrast, GraphST’s alignment performance degrades more significantly, leading to mismatches between different tissue regions.

Furthermore, our method achieves an average clustering performance improvement of over 56% compared to GraphST in this setting, demonstrating strong robustness against partial data loss. This advantage stems from S3RL’s graph-based alignment strategy, which ensures spatial continuity even when slice overlap is limited, thereby enabling more reliable clustering.

## III. Discussion

Spatial transcriptomics has revolutionized our understanding of tissue organization by providing spatially resolved gene expression profiles. However, existing analytical frameworks face persistent limitations in multimodal feature extraction and spatial representation learning. First, histological images often exhibit high structural homogeneity and blurred region boundaries, making it difficult for conventional models to extract meaningful semantic information. As a result, the contribution of image features to downstream analysis remains limited. Second, most graph construction strategies rely solely on spatial proximity (e.g., Euclidean distance or k-nearest neighbors), overlooking long-range but functionally similar spot relationships, while potentially introducing noisy connections between spatially adjacent yet biologically distinct spots. Lastly, current methods often lack constraints on representation separability in the latent space, leading to prototype collapse and indistinct cluster boundaries, which limits interpretability and robustness in downstream biological analysis.

In this study, we propose a separable spatial transcriptomics data analysis framework, S3RL, which integrates graph neural networks (GNNs) and hyperspherical prototype clustering to effectively analyze the organizational structure, gene expression patterns, and cross-slice alignment in spatial transcriptomics data. By constructing a graph structure based on spatial proximity and gene expression similarity, S3RL incorporates high-level semantic features from histological images, enabling more refined cell clustering and spatial structure recognition during the learning process. Unlike conventional methods, S3RL is not merely a graph-based modeling approach or a data enhancement technique; rather, it employs dynamic prototype learning and constraint mechanisms to ensure that high-dimensional expression features are more reasonably distributed in a low-dimensional space, thereby enhancing data stability and robustness. Moreover, S3RL adopts a hierarchical clustering strategy for spatial alignment and integration, balancing local cell-type resolution with global tissue structure alignment, thereby excelling in multi-scale spatial information fusion.

We validated the effectiveness of S3RL on several widely used spatial transcriptomics datasets, including DLPFC, Nanostring, Stereo-seq, and Slide-seqv2. Experimental results demonstrate that S3RL consistently achieves higher Adjusted Rand Index (ARI) scores across multiple datasets, outperforming state-of-the-art spatial transcriptomics analysis methods, such as GraphST, STAGATE, and BayesSpace, with an average improvement of over 40%. In the DLPFC dataset, S3RL exhibits enhanced cortical layer differentiation and achieves more accurate cell-type estimation in deconvolution analysis, highlighting its applicability in neural tissue research. Furthermore, in the Nanostring lung cancer dataset, we explored the performance of S3RL in gene expression enhancement, cell-cell communication analysis, and functional pathway enrichment. Results indicate that S3RL significantly improves the correlation between spatial transcriptomics data and TCGA bulk RNA-seq data, thereby enhancing the biological validity of spatial data and further demonstrating the potential of our method in cancer research. Additionally, in the cross-slice alignment task, S3RL effectively optimizes the integration of spatially aligned data, improving both data fusion and clustering performance. Even in cases where tissue regions exhibit only partial spatial overlap, S3RL maintains high clustering accuracy, making it particularly valuable for multi-tissue and multi-sample analyses in real-world applications.

Despite the outstanding performance of S3RL across multiple tasks, there remain several aspects that warrant further optimization. First, although this study enhances graph construction using prior visual features, the utilization of histological images still has room for improvement. Future research could incorporate self-supervised learning or multimodal contrastive learning strategies to ensure better generalization of image-derived features across different datasets. Second, while S3RL demonstrates strong adaptability in cross-slice alignment tasks, it may still be affected by data sparsity issues in low-signal, low-coverage spatial transcriptomics datasets, such as Slide-seq. This limitation could be addressed by integrating graph contrastive learning to further refine the model’s representations. Lastly, designing an end-to-end strategy that effectively balances clustering performance and multi-data integration remains a promising research direction.

Overall, S3RL provides an efficient, accurate, and biologically meaningful computational framework for spatial transcriptomics data analysis. In the future, it can be further extended to spatial multi-omics applications, facilitating broader biomedical research and applications across diverse biological contexts.

## IV. Method

The proposed S3RL framework consists of three main components: Semantic Information Extraction Module, Graph Construction Module, and Separable Representation Learning Module. We first introduce the data preprocessing and notations, followed by detailed descriptions of each module.

### A. Data Preprocessing and Notations

Spatial transcriptomics data generated by different platforms, including the NanoString CosMxTM SMI lung cancer dataset (Lung-91), Vizgen MERSCOPE mouse liver dataset L1R1, and 10x Visium datasets from the human dorsolateral prefrontal cortex (DLPFC), are preprocessed and represented in uniform format including gene expression matrix, spatial location, and histological images. For simplicity, the gene expression of spatial transcriptomics data can be characterized by **X** = {***x***_1_, …, ***x***_*n*_} ∈ ℝ^*n×m*^, the spatial location by **S** = {***s***_1_, …, ***s***_*n*_} ∈ ℝ^*n×*2^, and the histological image by **P** = {***p***_1_, …, ***p***_*n*_}, where *n* is the number of spots, *m* is the number of selected genes. As an unsupervised method, S3RL aims to learn the latent representation **Z** = {***z***_1_, …, ***z***_*n*_} ∈ ℝ^*n×d*^ of spots and simultaneously identify the labels of different spots ***l*** = {*l*_1_, …, *l*_*n*_} in an end-to-end manner, where *d* is the dimension of latent representation. Suppose there are *c* expected cell types or spatial domains. For the *i*-th spot from the Visium platform, a square patch centered on its location is regarded as its image with a size of 16 × 16 pixels, and for the *i*-th spot from the Nanostring CosMx platform, a square patch centered on its location is regarded as its image with a size of 60×60 pixels. We denote the graph by 𝒢 = (𝒱, ℰ), where 𝒱 is the set of nodes representing spots and ℰ is the set of edges connecting spots. Besides, we denote the adjacency matrix of 𝒢 by **A** ∈ ℝ^*n×n*^, where *a*_*ij*_ = 1 if (*i, j*) ∈ ℰ, otherwise *a*_*ij*_ = 0.

### B. Semantic Information Extraction Module

This module aims to extract high-level semantic information from histological images. To this end, we employ the SimCLR framework to learn the visual features of spots by maximizing the agreement between various augmented views of the same spot in a contrastive learning manner. For the *i*-th spot with image ***p***_*i*_, two augmented views 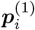 and 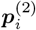 are regarded as a positive pair, and the augmented views of different images are regarded as negative pairs. Then we adopt the well-trained ResNet-50 network followed by simply Multilayer Perceptron (MLP) as the encoder ***f*** (•) to extract the visual features 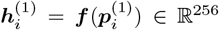 and 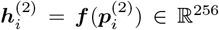. Contrastive learning aims to maximize the similarity between positive pairs and minimize the similarity between negative pairs, which can be formulated as the following loss function:

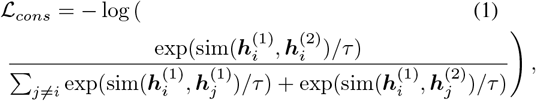

where sim(•, •) represents the cosine similarity between two features and *τ* is the temperature parameter. In this work, the random augmentations include random cropping, random flipping, and random color distortion. The learning rate is set to 0.005, *τ* is set to 0.5, the weight decay is set to 1*e*^−6^, and the model is trained for 500 epochs with the Adam optimizer and the batch size of 256. After training, we can obtain the visual features **H** = {***h***_1_, …, ***h***_*n*_} ∈ ℝ^*n×*256^ of *n* spots, encoding the high-level semantic information of histological images.

### C. Graph Construction Module

This module first converts the spatial location **S** into an undirected graph 𝒢= (𝒱,ℰ). Each spot in the graph 𝒢 connects to their *k* nearest neighbors such that ℰ = {(*i, j*)|*j* 𝒩_*i*_, where 𝒩_*i*_ denotes the set of *k* nearest neighbors of the *i*-th spot. Note that the neighboring nodes 𝒩_*i*_ can be determined by different distance or similarity metrics, and we adopt the commonest Euclidean distance in this work. Besides, this module further constructs a signed similarity graph based on the visual features to extract reliable pairwise relations between spots. Specifically, each spot is linked to another spot with a positive edge if they have the highest similarity and a negative edge if they have the lowest similarity based on the visual features. We denote the positive and negative edges by 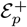 and 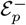, respectively. Similarly, we extract the prior knowledge inherent in gene profiles **X**. We calculate the pairwise similarity between spots based on the expression profiles **X** and retain the positive and negative edges according to the highest and lowest similarity, which are represented by 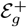 and 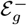, respectively.

The spatial graph 𝒢 presents local spatial information where spatially neighboring spots are more likely of the same cell type, but at the same time, it is incapable of capturing the global structure and high-level semantic information. Thus we introduce the positive and negative edges from the visual features and gene expression profiles to enhance the spatial graph. The positive edges indicate the possible homogeneity between spots, while the negative edges indicate the possible heterogeneity. We integrate the positive edges into the spatial graph such that the semantic graph 𝒢^*′*^ = (𝒱, ℰ^*′*^) and 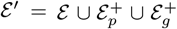. In this way, the graph 𝒢^*′*^ contains the spatial information and the semantic information from histological images and gene expression profiles, which facilitates the identification of distinct cell types and separating heterogeneous domains. Besides, the negative graph 𝒢^−^ = (𝒱, ℰ^−^), where 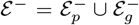, indicates dissimilarity of spots and is used to push them apart in the embedding space, for which we will explain more detailed in the following section.

### D. Separable Representation Learning

S3RL employs the popular GNNs as the encoder and decoder to learn informative low-dimension representations of spots by collaboratively taking into account the spatial information, the high-level semantic information, and the gene expression profiles. The encoder 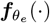 parameterized by *θ*_*e*_ takes the enhanced semantic graph 𝒢^*′*^ and the gene expression profiles **X** as input and outputs the low-dimension representation **Z**, which is followed by the symmetrical decoder 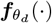 parameterized by *θ*_*d*_ to reconstruct the gene expression profiles. The encoder 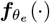 and decoder 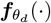 both consist of two-layer Graph Transformer Networks (GTNs) [58]. Formally, the (*l*+1)-th layer of the encoder is formulated as:

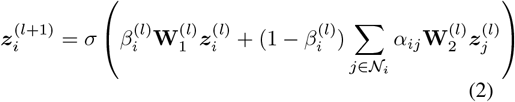

in which 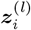 is the representation of the *i*-th spot in the *l*-th layer, 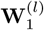 and 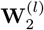 are the learnable transformation matrix. *σ*(•) is the activation function and S3RL adopts the ELU function. *α*_*ij*_ is the attention weight calculated by the scaled dot-product attention mechanism, and 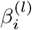 is a gated residual connection that balances the importance between neighboring nodes’ features and the *i*-th node’s own feature.

The symmetrical decoder aims to reconstruct the gene expression profiles in the same way as depicted in Eq. (2). Supposing the reconstructed profiles are 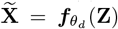, S3RL employs the Consine Error (CE) loss to measure the difference between the input and the reconstructed profiles as follows:

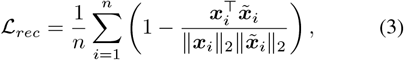

where ∥·∥_2_ denotes the ℓ_2_-norm of vectors.

Besides, the learned representation **Z** should maintain the topological structure of graphs 𝒢^*′*^ and 𝒢^−^, such that the spatial and semantic information is maximally preserved. S3RL introduces the next loss function to achieve it:

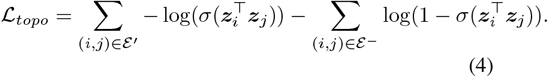

Minimizing ℒ_*topo*_ encourages spatially neighboring spots and semantically similar spots to be close in the embedding space while pushing dissimilar spots apart. In the training stage, S3RL randomly masks partial spots and replaces their profiles with a shard and learnable token to avoid overfitting and make the learned representation more expressive.

With the learned representations, one can apply a variety of off-the-shelf clustering algorithms to identify the cell type of each spot, such as *k*-means [59], Leiden [60], and Mclust [61], which are commonly used in many spatial transcriptomics studies [14], [18], [19]. However, this two-stage learning fashion, in which representation learning and clustering are performed separately, will lead to error propagation and information loss, resulting in suboptimal performance. Thus, S3RL proposes the hyperspherical prototype learning submodule to obtain the clustering results in an end-to-end manner.

S3RL first defines *c* prototypes {***p***_1_, ***p***_2_, …, ***p***_*c*_} on the *d*-dimension unit-hypersphere space such that the pairwise distance between any two prototypes is as large as possible. More importantly, each prototype is regarded as a centroid of a cell type and they are evenly scattered in the unit-hypersphere space. It can be achieved by minimizing the following problem using the gradient descent algorithm:

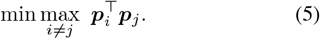

Then, S3RL maps the representation ***z***_*i*_ of the *i*-th spot into the same unit-hypersphere space as:

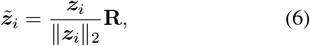

where **R** ∈ ℝ^*d×d*^ is a learnable orthogonal rotation matrix to align representations with *c* prototypes. In this way, the cell type affinity of the *i*-th spot with respect to the *j*-th prototypes can be calculated as:

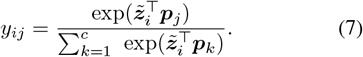

S3RL can obtain the final clustering label by assigning the *i*-th spot to the prototypes with the largest similarity as:

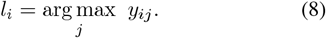

S3RL further introduces a hyperspherical regularization to promote the separability of the learned representations:

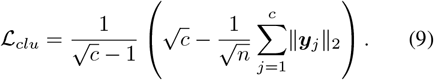

ℒ_*clu*_ encourages each spot to associate with a single prototype with higher confidence, while promoting separation between different types of spots, which improves the final clustering performance. Moreover, it ensures an even distribution of spots across prototypes, preventing empty clusters.

Instead of keeping the prototypes unchanged, prototypes are dynamically updated to adapt to diverse histology. The ideal situation derived by minimizing ℒ_*clu*_ is that each prototype is crowded with 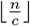 spots, *i*.*e*., for *j*-th prototype, the vector 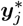 has 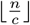 ones and 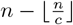 zeros. Then, S3RL can measure the difference between the current clustering results with the ideal one as follows:

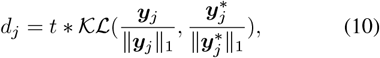

where 𝒦ℒ(•) denotes the Kullback-Leibler divergence and it can be replaced by other metrics, such as Spearman correlation, Jensen-Shannon divergence, and so on. *t* is a hyperparameter to adjust the scale of *d*_*j*_, and ∥ • ∥_1_ denotes the ℓ_1_-norm of vectors. The coefficient *d*_*j*_ reflects the degree of spots assigned to the *j*-th prototype deviating from the ideal one. Thus, S3RL can update the *j*-th prototype to minimize the deviation as:

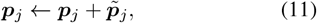

where 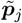 is sampled from the guassian distribution 𝒩 (**0**, *d*_*j*_**I**) with the mean of **0** and the variance of *d*_*j*_**I**. S3RL employs the reparameterization trick maintain the differentiability of the prototype update process.

The overall loss function of this separable representation learning module consists of the reconstruction loss, the topological loss, and the clustering loss:

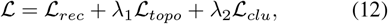

where *λ*_1_ and *λ*_2_ are hyperparameters to balance the importance of different losses. The model is trained end-to-end by minimizing the loss function ℒ using the Adam optimizer, for at most 2000 epochs.

### E. Multi-Slice Alignment and Joint Training

In multi-slice spatial transcriptomics analysis, balancing clustering performance with alignment and data integration remains a crucial challenge. While S3RL demonstrates outstanding performance in cell clustering, there exists an inherent trade-off between achieving optimal alignment across slices and preserving robust clustering results. To mitigate this conflict, we employ a spatial translation-only alignment strategy, avoiding distortions that may artificially enhance alignment at the cost of reduced clustering performance.

Specifically, we leverage PASTE2, a state-of-the-art partial alignment method, to perform global and local multi-slice alignment. PASTE2 optimally registers **only the overlapping regions between slices using a weighted Gromo–Wasserstein optimal transport strategy, thereby reducing interference from non-overlapping areas and enhancing robustness. We adopt the default parameter settings and use the `partial_pairwise_align` function to align the slices. For complete slices and partially cropped slices, we set the *s* parameter to 1 and 0.7, respectively, ensuring that the alignment remains effective under different levels of tissue overlap.

Once all slices are aligned to a common reference coordinate system, we merge them into a unified dataset for joint representation learning with S3RL. This enables similar spots to leverage neighboring spot expression patterns more effectively, ensuring both spatial coherence and feature consistency. As a result, our approach not only enhances the integration of multiple spatial slices but also maintains superior clustering accuracy, setting a robust foundation for downstream spatial transcriptomics analysis.

### F. Ligand-Receptor Pair Construction and Evaluation Metrics

To evaluate the effectiveness of different enhancement methods in ligand-receptor (L-R) identification, we first constructed a reliable L-R interaction set by integrating data from the IUPHAR/BPS Guide to Pharmacology [46] and the SiGra-aggregated database [20]. After redundancy removal, we curated 2,864 unique L-R pairs, among which 480 (189 ligands and 159 receptors) were present in the NanoString CosMx dataset. Using CellChat, we calculated the significance of each L-R interaction across cell clusters based on −log_10_(FDR) and visualized the results in a scatter plot, comparing raw and enhanced data. In the scatter plot (Supplementary Fig. S15 top), the x-axis represents raw significance, and the y-axis represents significance after enhancement. Each point corresponds to a specific L-R pair: the top-left quadrant represents L-R pairs that are newly significant only after enhancement, the bottom-right quadrant shows those that lost significance after enhancement, the top-right quadrant indicates consistently significant L-R pairs, and the bottom-left quadrant marks pairs that remain non-significant in both conditions.

To further quantify the model performance in L-R identification, we defined three complementary metrics.

The *Enhanced Specificity Ratio (ESR)* is defined as 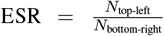, reflecting the method’s ability to discover new L-R pairs not detected in the raw data relative to those lost after enhancement. The *Effective Enhancement Percentage (EEP)* is defined as 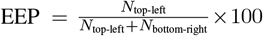, indicating the proportion of newly discovered significant L-R pairs among all changed ones. The *Specificity Stability Percentage (SSP)* is computed as 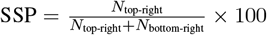, measuring how well the method preserves originally significant L-R pairs during enhancement.

Here, *N*_top-left_ denotes the number of L-R pairs that become significant only after enhancement, *N*_top-right_ refers to those consistently significant before and after enhancement, and *N*_bottom-right_ represents L-R pairs that were significant in raw data but lost significance post-enhancement. Together, these metrics offer a comprehensive and interpretable assessment of a model’s capacity to enhance biologically meaningful cell-cell communication signals while minimizing disruption to existing ones.

### G. Benchmarking Methods and Performance Evaluation

To evaluate the performance of our proposed model, we conducted a comprehensive comparison against several state-of-the-art methods, including BayesSpace, Giotto, Seurat, SEDR, SiGra, conST, SpaceFlow, spaGCN, and STAGATE, using Adjusted Rand Index (ARI) as the evaluation metric (Figure 2). The ARI measures the alignment between predicted spatial clusters and the ground truth annotations, providing an indication of how well each method captures the underlying biological structure of the tissue samples.

Our model consistently outperforms the competing methods across multiple datasets, including lung cancer, mouse brain anterior, and human breast cancer, demonstrating superior accuracy in segmenting distinct spatial regions. Specifically, our method achieves a significantly higher ARI score, indicating that it better preserves the spatial and biological context of the tissue. The enhanced representation of gene expression and spatial structure in our model allows for more accurate delineation of complex tissue microenvironments compared to existing methods.

The benchmarking results further confirm the robustness of our model across various tissue types and biological conditions, providing strong evidence of its generalizability and effectiveness in spatial transcriptomics analysis.

### H. Identifying Differentially Expressed Genes and Adjacent Cell Communications

In addition to clustering performance, we also explored the identification of differentially expressed genes (DEGs) and the analysis of adjacent cell communications. For this, we used the Wilcoxon test to identify DEGs in each spatial region, comparing gene expression between neighboring cell clusters. We set a threshold of log2 fold change greater than 1 and adjusted p-value less than 0.05 for significant DEGs.

Furthermore, we analyzed cell-cell communication by calculating the interaction strength between adjacent cell pairs. This was done by aggregating the ligand-receptor interaction scores for neighboring cells and normalizing the interaction strengths based on the expression levels of key signaling molecules. The communication strength between adjacent cell pairs reflects the level of molecular crosstalk, which is critical in understanding the role of different cell types in the tissue microenvironment.

The results revealed several important communication pathways, including growth factor signaling, immune cell interactions, and tumor-stroma crosstalk. Notably, enhanced data from our model provided a clearer view of these communication patterns, uncovering interactions that were less visible in the raw data. This enhanced resolution of cell-cell communication networks highlights the biological relevance of our model and its ability to capture key molecular interactions in spatial transcriptomics datasets.

To validate the effectiveness of the enhanced count matrix, we analyzed both the original and enhanced matrices using Seurat and CellChat. Firstly, we employed Seurat’s Wilcoxon Rank Sum test to identify marker genes, setting the significance level at 0.05. Following the identification of marker genes, we selected upregulated genes with log_2_ FC ≥ 0.2 and adjusted *p*-value ≤ 0.05, as well as downregulated genes with log_2_ FC ≤− 0.25 and adjusted *p*-value ≤ 0.05, then plotted a volcano plot.

Subsequently, utilizing the CellChatDB database of human ligand-receptor interactions, we modeled ligand-receptor mediated signaling interactions based on the law of mass action. This approach enabled us to determine the communication probabilities between all cell groups for each ligand-receptor pair. We then calculated the communication probabilities between each cell type by weighting the probabilities of the relevant ligand-receptor pairs, and plotted ligand-receptor heatmaps and communication strength plots between different cell types.

We then compared the ligand-receptor networks and marker genes of the enhanced and original matrices, identifying newly emerged ligand-receptor pathways in the enhanced matrix. These corresponding genes were highlighted in the marker genes volcano plot. Additionally, we grouped the data by cell type and used the count matrix to plot violin plots of the gene intensities for the highlighted genes in the marker genes volcano plot.

## V. Statistics and reproducibility

No statistical method was used to predetermine the sample size. No data were excluded from the analyses. The experiments were not randomized. The investigators were not blinded to allocation during experiments and outcome assessment.

## VI. Reporting summary

Further information on research design is available in the Nature Portfolio Reporting Summary linked to this article.

## VII. Data availability

All datasets used in this study are publicly available and widely adopted as benchmarks for evaluating spatial transcriptomics methods. The 10X Visium spatial transcriptomics dataset of the human dorsolateral prefrontal cortex (DLPFC), comprising 12 tissue slices with manually annotated cortical layers, was obtained from Maynard et al. [24]. The Mouse Brain Anterior and Human Breast Cancer datasets were downloaded from the official 10X Genomics website (https://www.10xgenomics.com/resources/datasets), both sequenced using the Visium platform. To assess the model’s robustness on large-scale pathological tissues, we utilized the Nanostring CosMx Spatial Molecular Imaging (SMI) dataset, which includes 20 formalin-fixed, paraffin-embedded (FFPE) lung cancer slices with expert-labeled cell type annotations. This dataset is publicly available via the Nanostring official website (https://nanostring.com/products/cosmx-spatial-molecular-imager/nsclc-ffpe-dataset/). Additionally, we included high-resolution mouse olfactory bulb Stereo-seq data and mouse hippocampus Slide-seqV2 data from their original publications [5], [7] to validate performance in subcellular contexts. For the integration analysis of scRNA-seq and spatial transcriptomics (ST), we used a single-nucleus RNA-seq (snRNA-seq) dataset derived from the human dorsolateral prefrontal cortex (BA9), comprising 78,886 nuclei and 30,062 genes, obtained from the GEO database (accession number GSE144136) [29]. The processed data used in this study have been uploaded to Zenodo and are freely available at: https://zenodo.org/records/15222342.

## VIII. Code availability

An open-source Python implementation of the proposed method is available at: https://github.com/AI4Bread/S3RL.

## IX. ACKNOWLEDGEMENTS

This research was supported by National Natural Science Foundation of China (Grant No. 62303372 to L.F., & No.GYKP034 to H.S.), Zhejiang Provincial Natural Science Foundation of China under Grant No. LQ23F020018 to L.F., Young Talent Fund of Xi’an Association for Science and Technology (No. 959202313033 to L.F.), Project funded by China Post-doctoral Science Foundation (No. 2023M742794 to L.F.), and Postdoctoral Research Project in Shaanxi Province (2023BSHEDZZ34 to L.F.), Fundamental Research Funds for the Central Universities (xzy012024091 to L.F.).

## X. Author contributions

D.W. and L.F. conceptualized and supervised the study, L.F., P.W., G.X., J.L., H.S. and D.W. performed model design, data analysis, software development, and manuscript writing, H.S. supervised the study. All the authors performed manuscript writing.

## XI. Competing interests

The authors declare no competing interests.

## XII. Inclusion & Ethics Statement

This research does not involve any ethical concerns related to resource-poor settings, nor does it present any risks or harm to the participants.

## XIII. Additional information

**Supplementary information** The online version contains supplementary material available at.

**Correspondence and requests for materials** should be addressed to.

## References

[1] P. L. Ståhl, F. Salmén, S. Vickovic, A. Lundmark, J. F. Navarro, J. Magnusson, S. Giacomello, M. Asp, J. O. Westholm, M. Huss et al., “Visualization and analysis of gene expression in tissue sections by spatial transcriptomics,” Science, vol. 353, no. 6294, pp. 78–82, 2016.

[2] P. V. Kharchenko, “The triumphs and limitations of computational methods for scrna-seq,” Nature methods, vol. 18, no. 7, pp. 723–732, 2021.

[3] 10x Genomics, “10x genomics: Visium spatial gene expression,” 2024. [Online]. Available: https://www.10xgenomics.com/

[4] S. G. Rodriques, R. R. Stickels, A. Goeva, C. A. Martin, E. Murray, C. R. Vanderburg, J. Welch, L. M. Chen, F. Chen, and E. Z. Macosko, “Slide-seq: A scalable technology for measuring genome-wide expression at high spatial resolution,” Science, vol. 363, no. 6434, pp. 1463–1467, 2019.

[5] R. R. Stickels, E. Murray, P. Kumar, J. Li, J. L. Marshall, D. J. Di Bella, P. Arlotta, E. Z. Macosko, and F. Chen, “Highly sensitive spatial transcriptomics at near-cellular resolution with slide-seqv2,” Nature biotechnology, vol. 39, no. 3, pp. 313–319, 2021.

[6] Y. Liu, M. Yang, Y. Deng, G. Su, A. Enninful, C. C. Guo, T. Tebaldi, D. Zhang, D. Kim, Z. Bai et al., “High-spatial-resolution multi-omics sequencing via deterministic barcoding in tissue,” Cell, vol. 183, no. 6, pp. 1665–1681, 2020.

[7] A. Chen, S. Liao, M. Cheng, K. Ma, L. Wu, Y. Lai, X. Qiu, J. Yang, J. Xu, S. Hao et al., “Spatiotemporal transcriptomic atlas of mouse organogenesis using dna nanoball-patterned arrays,” Cell, vol. 185, no. 10, pp. 1777–1792, 2022.

[8] R. Zahedi, R. Ghamsari, A. Argha, C. Macphillamy, A. Beheshti, R. Alizadehsani, N. H. Lovell, M. Lotfollahi, and H. Alinejad-Rokny, “Deep learning in spatially resolved transcriptomics: a comprehensive technical view,” Briefings in Bioinformatics, vol. 25, no. 2, p. bbae082, 2024.

[9] E. Lubeck, A. F. Coskun, T. Zhiyentayev, M. Ahmad, and L. Cai, “Single-cell in situ rna profiling by sequential hybridization,” Nature methods, vol. 11, no. 4, pp. 360–361, 2014.

[10] Z. Yao, C. T. van Velthoven, M. Kunst, M. Zhang, D. McMillen, C. Lee, W. Jung, J. Goldy, A. Abdelhak, M. Aitken et al., “A high-resolution transcriptomic and spatial atlas of cell types in the whole mouse brain,” Nature, vol. 624, no. 7991, pp. 317–332, 2023.

[11] L. Fei, H. Chen, L. Ma, W. E R. Wang, X. Fang, Z. Zhou, H. Sun, J. Wang, M. Jiang et al., “Systematic identification of cell-fate regulatory programs using a single-cell atlas of mouse development,” Nature Genetics, vol. 54, no. 7, pp. 1051–1061, 2022.

[12] S. Jin, C. F. Guerrero-Juarez, L. Zhang, I. Chang, R. Ramos, C.-H. Kuan, P. Myung, M. V. Plikus, and Q. Nie, “Inference and analysis of cell-cell communication using cellchat,” Nature communications, vol. 12, no. 1, p. 1088, 2021.

[13] R. Dries, Q. Zhu, R. Dong, C.-H. L. Eng, H. Li, K. Liu, Y. Fu, T. Zhao, A. Sarkar, F. Bao et al., “Giotto: a toolbox for integrative analysis and visualization of spatial expression data,” Genome biology, vol. 22, pp. 1–31, 2021.

[14] J. Hu, X. Li, K. Coleman, A. Schroeder, N. Ma, D. J. Irwin, E. B. Lee, R. T. Shinohara, and M. Li, “Spagcn: Integrating gene expression, spatial location and histology to identify spatial domains and spatially variable genes by graph convolutional network,” Nature methods, vol. 18, no. 11, pp. 1342–1351, 2021.

[15] J. Li, S. Chen, X. Pan, Y. Yuan, and H.-B. Shen, “Cell clustering for spatial transcriptomics data with graph neural networks,” Nature Computational Science, vol. 2, no. 6, pp. 399–408, 2022.

[16] D. Pham, X. Tan, J. Xu, L. F. Grice, P. Y. Lam, A. Raghubar, J. Vukovic, M. J. Ruitenberg, and Q. Nguyen, “stlearn: integrating spatial location, tissue morphology and gene expression to find cell types, cell-cell interactions and spatial trajectories within undissociated tissues,” BioRxiv, pp. 2020–05, 2020.

[17] E. Zhao, M. R. Stone, X. Ren, J. Guenthoer, K. S. Smythe, T. Pulliam, S. R. Williams, C. R. Uytingco, S. E. Taylor, P. Nghiem et al., “Spatial transcriptomics at subspot resolution with bayesspace,” Nature biotechnology, vol. 39, no. 11, pp. 1375–1384, 2021.

[18] K. Dong and S. Zhang, “Deciphering spatial domains from spatially resolved transcriptomics with an adaptive graph attention auto-encoder,” Nature communications, vol. 13, no. 1, p. 1739, 2022.

[19] Y. Long, K. S. Ang, M. Li, K. L. K. Chong, R. Sethi, C. Zhong, H. Xu, Z. Ong, K. Sachaphibulkij, A. Chen et al., “Spatially informed clustering, integration, and deconvolution of spatial transcriptomics with graphst,” Nature Communications, vol. 14, no. 1, p. 1155, 2023.

[20] Z. Tang, Z. Li, T. Hou, T. Zhang, B. Yang, J. Su, and Q. Song, “Sigra: single-cell spatial elucidation through an image-augmented graph transformer,” Nature Communications, vol. 14, no. 1, p. 5618, 2023.

[21] H. Xu, H. Fu, Y. Long, K. S. Ang, R. Sethi, K. Chong, M. Li, R. Uddamvathanak, H. K. Lee, J. Ling et al., “Unsupervised spatially embedded deep representation of spatial transcriptomics,” Genome Medicine, vol. 16, no. 1, p. 12, 2024.

[22] T. Chen, S. Kornblith, M. Norouzi, and G. E. Hinton, “A simple framework for contrastive learning of visual representations,” in Proceedings of the 37th International Conference on Machine Learning, 2020, pp. 1597–1607.

[23] Z. Wu, S. Pan, F. Chen, G. Long, C. Zhang, and P. S. Yu, “A comprehensive survey on graph neural networks,” IEEE transactions on neural networks and learning systems, vol. 32, no. 1, pp. 4–24, 2020.

[24] K. R. Maynard, L. Collado-Torres, L. M. Weber, C. Uytingco, B. K. Barry, S. R. Williams, J. L. Catallini, M. N. Tran, Z. Besich, M. Tippani et al., “Transcriptome-scale spatial gene expression in the human dorsolateral prefrontal cortex,” Nature neuroscience, vol. 24, no. 3, pp. 425–436, 2021.

[25] Y. Hao, T. Stuart, M. H. Kowalski, S. Choudhary, P. Hoffman, A. Hartman, A. Srivastava, G. Molla, S. Madad, C. Fernandez-Granda et al., “Dictionary learning for integrative, multimodal and scalable single-cell analysis,” Nature biotechnology, vol. 42, no. 2, pp. 293–304, 2024.

[26] Y. Zong, T. Yu, X. Wang, Y. Wang, Z. Hu, and Y. Li, “const: an interpretable multi-modal contrastive learning framework for spatial transcriptomics,” BioRxiv, pp. 2022–01, 2022.

[27] H. Ren, B. L. Walker, Z. Cang, and Q. Nie, “Identifying multicellular spatiotemporal organization of cells with spaceflow,” Nature communications, vol. 13, no. 1, p. 4076, 2022.

[28] S. A. Mauney, C. Y. Pietersen, K.-C. Sonntag, and T.-U. W. Woo, “Differentiation of oligodendrocyte precursors is impaired in the prefrontal cortex in schizophrenia,” Schizophrenia research, vol. 169, no. 1-3, pp. 374–380, 2015.

[29] S. Bae, K. J. Na, J. Koh, D. S. Lee, H. Choi, and Y. T. Kim, “Celldart: cell type inference by domain adaptation of single-cell and spatial transcriptomic data,” Nucleic acids research, vol. 50, no. 10, pp. e57–e57, 2022.

[30] O. A. Bayraktar, T. Bartels, S. Holmqvist, V. Kleshchevnikov, A. Martirosyan, D. Polioudakis, L. Ben Haim, A. M. Young, M. Y. Batiuk, K. Prakash et al., “Astrocyte layers in the mammalian cerebral cortex revealed by a single-cell in situ transcriptomic map,” Nature neuroscience, vol. 23, no. 4, pp. 500–509, 2020.

[31] S. He, R. Bhatt, C. Brown, E. A. Brown, D. L. Buhr, K. Chantranuvatana, P. Danaher, D. Dunaway, R. G. Garrison, G. Geiss et al., “High-plex imaging of rna and proteins at sub-cellular resolution in fixed tissue by spatial molecular imaging,” Nature biotechnology, vol. 40, no. 12, pp. 1794–1806, 2022.

[32] D. Li, X. Guo, K. Yang, Y. Yang, W. Zhou, Y. Huang, X. Liang, J. Su, L. Jiang, J. Li et al., “Epcam-targeting car-t cell immunotherapy is safe and efficacious for epithelial tumors,” Science Advances, vol. 9, no. 48, p. eadg9721, 2023.

[33] X. Shi, W. Liang, W. Yang, R. Xia, and Y. Song, “Decorin is responsible for progression of non-small-cell lung cancer by promoting cell proliferation and metastasis,” Tumor Biology, vol. 36, pp. 3345–3354, 2015.

[34] T. Wen, X. Zhang, Y. Gao, H. Tian, L. Fan, and P. Yang, “Sox4-bmi1 axis promotes non-small cell lung cancer progression and facilitates angiogenesis by suppressing znf24,” Cell Death & Disease, vol. 15, no. 9, p. 698, 2024.

[35] Y. Xie, N. Su, J. Yang, Q. Tan, S. Huang, M. Jin, Z. Ni, B. Zhang, D. Zhang, F. Luo et al., “Fgf/fgfr signaling in health and disease,” Signal transduction and targeted therapy, vol. 5, no. 1, p. 181, 2020.

[36] P. Zhang, L. Yue, Q. Leng, C. Chang, C. Gan, T. Ye, and D. Cao, “Targeting fgfr for cancer therapy,” Journal of Hematology & Oncology, vol. 17, no. 1, p. 39, 2024.

[37] V. Yellapragada, N. Eskici, Y. Wang, S. Madhusudan, K. Vaaralahti, T. Tuuri, and T. Raivio, “Fgf8–fgfr1 signaling regulates human gnrh neuron differentiation in a time-and dose-dependent manner,” Disease Models & Mechanisms, vol. 15, no. 8, p. dmm049436, 2022.

[38] X. Ke, S. Xia, W. Yu, S. Mabry, Q. Fu, H. L. Menden, V. Sampath, and R. H. Lane, “Delta like 4 regulates cerebrovascular development and endothelial integrity via dll4-notch-cldn5 pathway and is vulnerable to neonatal hyperoxia,” The Journal of physiology, vol. 602, no. 10, pp. 2265–2285, 2024.

[39] J.-f. Zhang, Y. Chen, X.-x. Qiu, W.-l. Tang, J.-d. Zhang, J.-h. Huang, G.-s. Lin, X.-f. Wang, and Z.-x. Lin, “The vascular deltalike ligand-4 (dll4)-notch4 signaling correlates with angiogenesis in primary glioblastoma: an immunohistochemical study,” Tumor Biology, vol. 37, pp. 3797–3805, 2016.

[40] P. Nowak-Sliwinska, J. R. van Beijnum, E. J. Huijbers, P. C. Gasull, L. Mans, A. Bex, and A. W. Griffioen, “Oncofoetal insulin receptor isoform a marks the tumour endothelium; an underestimated pathway during tumour angiogenesis and angiostatic treatment,” British journal of cancer, vol. 120, no. 2, pp. 218–228, 2019.

[41] H. R. Ferguson, M. P. Smith, and C. Francavilla, “Fibroblast growth factor receptors (fgfrs) and noncanonical partners in cancer signaling,” Cells, vol. 10, no. 5, p. 1201, 2021.

[42] M. A. Krook, J. W. Reeser, G. Ernst, H. Barker, M. Wilberding, G. Li, H.-Z. Chen, and S. Roychowdhury, “Fibroblast growth factor receptors in cancer: genetic alterations, diagnostics, therapeutic targets and mechanisms of resistance,” British Journal of Cancer, vol. 124, no. 5, pp. 880–892, 2021.

[43] E. B. Pasquale, “Eph receptors and ephrins in cancer progression,” Nature Reviews Cancer, vol. 24, no. 1, pp. 5–27, 2024.

[44] G. Hanover, F. S. Vizeacoumar, S. L. Banerjee, R. Nair, R. Dahiya, A. I. Osornio-Hernandez, A. M. Morales, T. Freywald, J. P. Himanen, B. M. Toosi et al., “Integration of cancer-related genetic landscape of eph receptors and ephrins with proteomics identifies a crosstalk between ephb6 and egfr,” Cell reports, vol. 42, no. 7, 2023.

[45] B. D. Ferguson, R. Liu, C. E. Rolle, Y.-H. C. Tan, V. Krasnoperov, R. Kanteti, M. S. Tretiakova, G. M. Cervantes, R. Hasina, R. D. Hseu et al., “The ephb4 receptor tyrosine kinase promotes lung cancer growth: a potential novel therapeutic target,” PloS one, vol. 8, no. 7, p. e67668, 2013.

[46] J. L. Sharman, H. E. Benson, A. J. Pawson, V. Lukito, C. P. Mpamhanga, V. Bombail, A. P. Davenport, J. A. Peters, M. Spedding, A. J. Harmar et al., “Iuphar-db: updated database content and new features,” Nucleic acids research, vol. 41, no. D1, pp. D1083–D1088, 2012.

[47] T. G. Araújo, S. T. S. Mota, H. S. V. Ferreira, M. A. Ribeiro, L. R. Goulart, and L. Vecchi, “Annexin a1 as a regulator of immune response in cancer,” Cells, vol. 10, no. 9, p. 2245, 2021.

[48] K. Shien, V. A. Papadimitrakopoulou, D. Ruder, C. Behrens, L. Shen, N. Kalhor, J. Song, J. J. Lee, J. Wang, X. Tang et al., “Jak1/stat3 activation through a proinflammatory cytokine pathway leads to resistance to molecularly targeted therapy in non–small cell lung cancer,” Molecular cancer therapeutics, vol. 16, no. 10, pp. 2234–2245, 2017.

[49] C. Grimm, S. Frässle, C. Steger, L. von Ziegler, O. Sturman, N. Shemesh, D. Peleg-Raibstein, D. Burdakov, J. Bohacek, K. E. Stephan et al., “Optogenetic activation of striatal d1r and d2r cells differentially engages downstream connected areas beyond the basal ganglia,” Cell reports, vol. 37, no. 13, 2021.

[50] X.-L. Gu, Z.-L. Ou, F.-J. Lin, X.-L. Yang, J.-M. Luo, Z.-Z. Shen, and Z.-M. Shao, “Expression of cxcl14 and its anticancer role in breast cancer,” Breast cancer research and treatment, vol. 135, pp. 725–735, 2012.

[51] X. Zhong, W. Zhang, and T. Sun, “Ddr1 promotes breast tumor growth by suppressing antitumor immunity,” Oncology Reports, vol. 42, no. 6, pp. 2844–2854, 2019.

[52] S. Zeppilli, T. Ackels, R. Attey, N. Klimpert, K. D. Ritola, S. Boeing, A. Crombach, A. T. Schaefer, and A. Fleischmann, “Molecular characterization of projection neuron subtypes in the mouse olfactory bulb,” Elife, vol. 10, p. e65445, 2021.

[53] H. E. Marei, A.-E. Ahmed, F. Michetti, M. Pescatori, R. Pallini, P. Casalbore, C. Cenciarelli, and M. Elhadidy, “Gene expression profile of adult human olfactory bulb and embryonic neural stem cell suggests distinct signaling pathways and epigenetic control,” PLoS One, vol. 7, no. 4, p. e33542, 2012.

[54] L. R. Saraiva, X. Ibarra-Soria, M. Khan, M. Omura, A. Scialdone, P. Mombaerts, J. C. Marioni, and D. W. Logan, “Hierarchical deconstruction of mouse olfactory sensory neurons: from whole mucosa to single-cell rna-seq,” Scientific reports, vol. 5, no. 1, p. 18178, 2015.

[55] L. Loo, J. M. Simon, L. Xing, E. S. McCoy, J. K. Niehaus, J. Guo, E. Anton, and M. J. Zylka, “Single-cell transcriptomic analysis of mouse neocortical development,” Nature communications, vol. 10, no. 1, p. 134, 2019.

[56] K. Ikeda, T. Onaka, M. Yamakado, J. Nakai, T.-o. Ishikawa, M. M. Taketo, and K. Kawakami, “Degeneration of the amygdala/piriform cortex and enhanced fear/anxiety behaviors in sodium pump α2 subunit (atp1a2)-deficient mice,” Journal of Neuroscience, vol. 23, no. 11, pp. 4667–4676, 2003.

[57] P. S. Villar, R. Delgado, C. Vergara, J. G. Reyes, and J. Bacigalupo, “Energy requirements of odor transduction in the chemosensory cilia of olfactory sensory neurons rely on oxidative phosphorylation and glycolytic processing of extracellular glucose,” Journal of Neuroscience, vol. 37, no. 23, pp. 5736– 5743, 2017.

[58] S. Yun, M. Jeong, R. Kim, J. Kang, and H. J. Kim, “Graph transformer networks,” Advances in neural information processing systems, vol. 32, 2019.

[59] J. A. Hartigan and M. A. Wong, “Algorithm as 136: A k-means clustering algorithm,” Journal of the royal statistical society. series c (applied statistics), vol. 28, no. 1, pp. 100–108, 1979.

[60] V. A. Traag, L. Waltman, and N. J. Van Eck, “From louvain to leiden: guaranteeing well-connected communities,” Scientific reports, vol. 9, no. 1, p. 5233, 2019.

[61] L. Scrucca, M. Fop, T. B. Murphy, and A. E. Raftery, “mclust 5: clustering, classification and density estimation using gaussian finite mixture models,” The R journal, vol. 8, no. 1, p. 289, 2016.

