## Supplementary file for "Separable Spatial Single-cell Transcriptome Representation Learning via Graph Transformer and Hyperspherical Prototype Clustering"

#### 1 Latent Space Clustering Results on DPFLC dataset

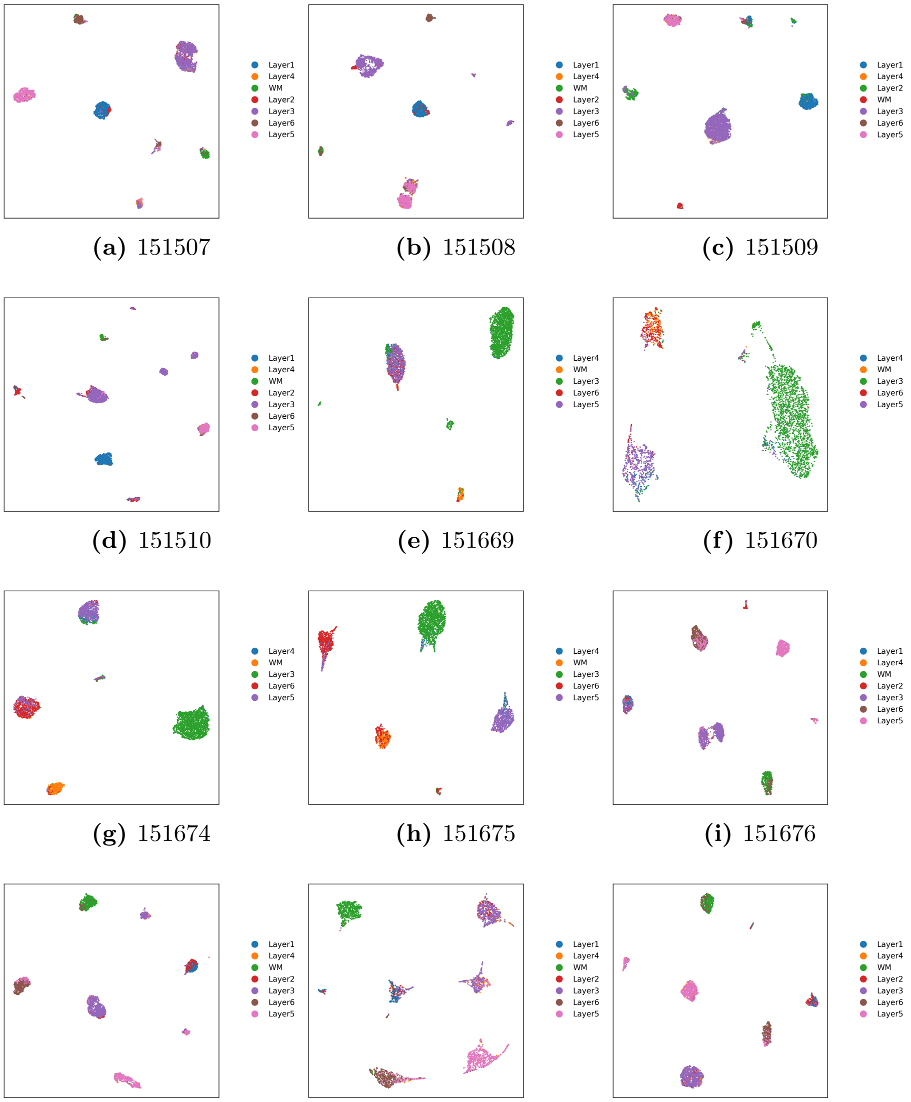

(a) 151507 (b) 151508 (c) 151509

(d) 151510 (e) 151669 (f) 151670

(g) 151674 (h) 151675 (i) 151676

(j) 151674 (k) 151675 (l) 151676

**Fig. S1: Clustering results in the latent space for 12 DLPFC slices from the 10X Visium dataset using S3RL.** Each subplot represents the latent space embedding of one slice, where spots are colored according to their predicted clusters.

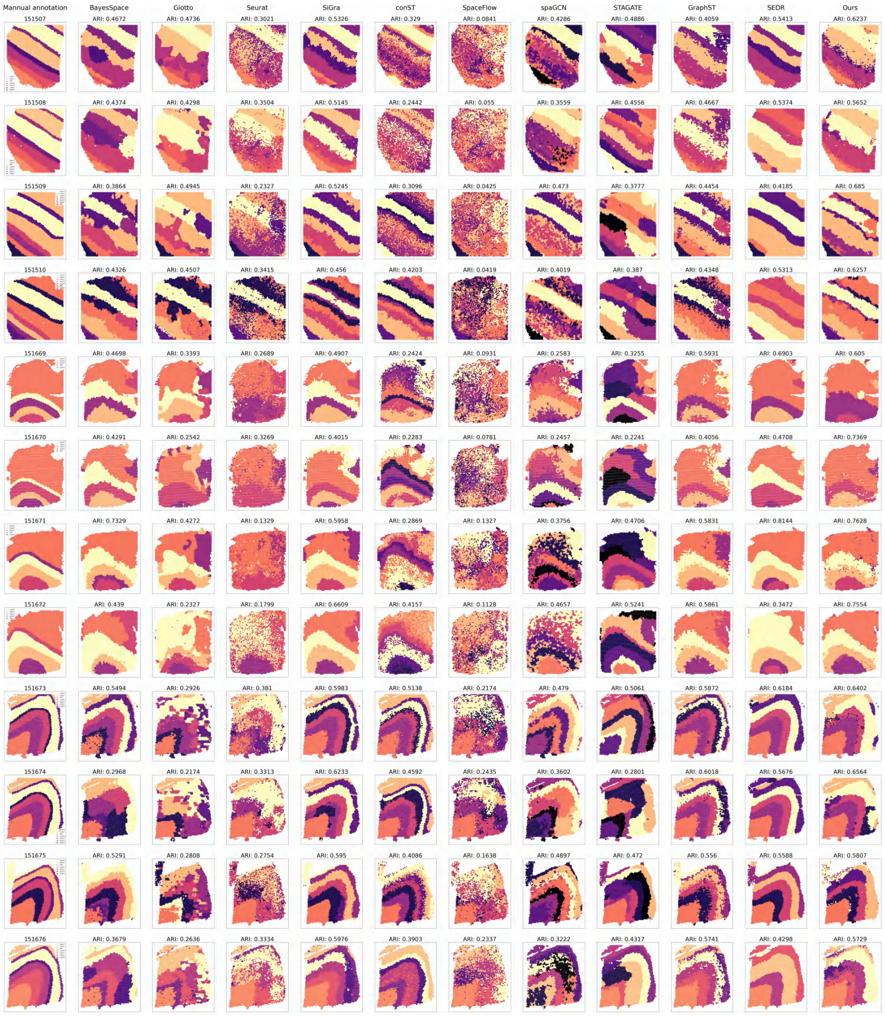

**Fig. S2: Visualization of clustering results across 12 DLPFC slices from the 10X Visium dataset.** The first column presents manual annotations as ground truth, while the remaining columns show clustering results obtained from various spatial transcriptomics analysis methods, including BayesSpace, Giotto, Seurat, SiGra, conST, SpaceFlow, spaGCN, STAGATE, GraphST, S3RL, and our proposed method. Each subplot represents a different tissue slice, with spots colored according to their predicted clusters. The Adjusted Rand Index (ARI) values are displayed for each method, indicating clustering performance relative to ground truth annotations.

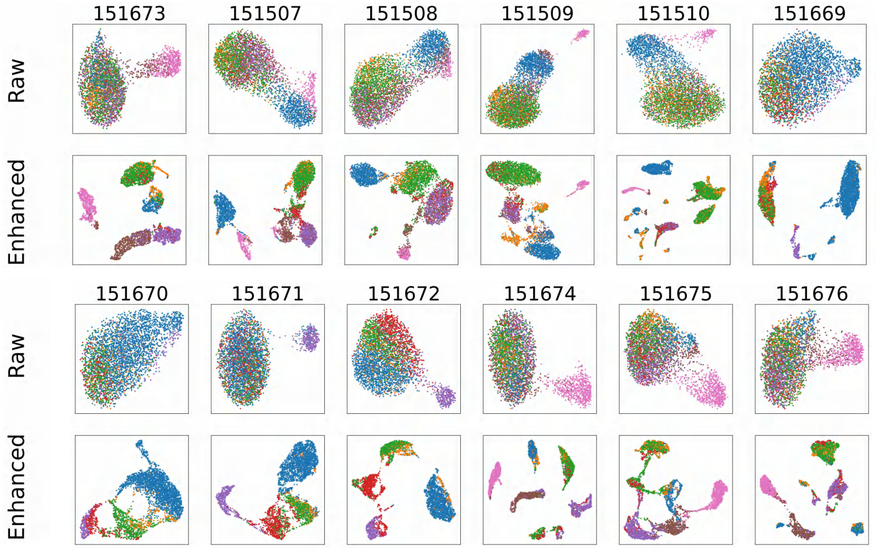

**Fig. S3: Raw and enhanced clustering performance visualization in the latent space of DLPFC slices.** The first and third rows represent the raw data, and the second and fourth rows correspond to the S3RL framework-enhanced data. The S3RL enhanced embeddings show exceptionally elevated separation and compactness compared to the raw embeddings, facilitating the identification of cellular heterogeneity within the DLPFC tissue architecture.

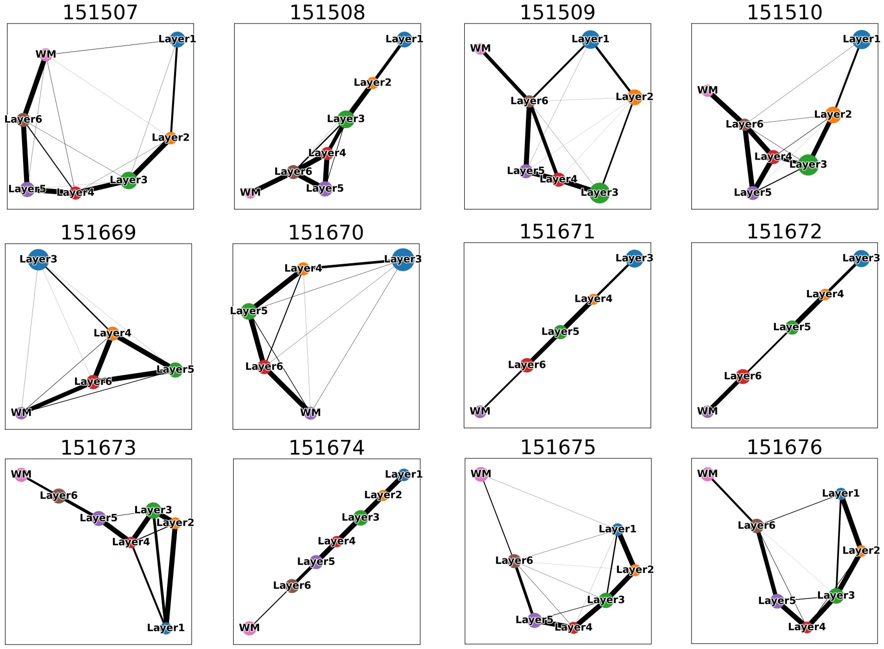

**Fig. S4: Cell trajectory analysis on 12 slices of the 10X Visium DLPFC dataset using S3RL.** Each subplot represents a different tissue slice, where layers (e.g., Layer1–Layer6 and WM) are connected based on inferred cellular transitions. The thickness of the edges indicates the strength of inferred connections, reflecting potential spatial interactions and lineage relationships among layers. This visualization demonstrates the ability of S3RL to preserve biologically meaningful spatial connectivity while distinguishing distinct cellular compartments.

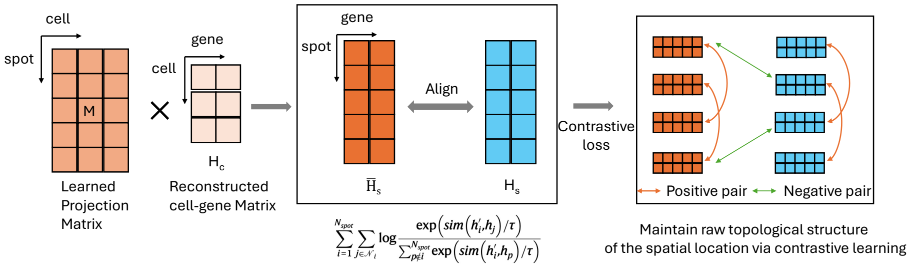

**Fig. S5: Overview of the contrastive learning framework for spatial gene expression alignment.** Instead of using an auto-encoder, the reconstructed cell-gene expression matrix $H_c$ is directly utilized. A learned projection matrix $M$ maps single-cell RNA-seq data to the spatial transcriptomics domain, generating the reconstructed spatial gene expression matrix $\overline{H}_s$. The alignment process ensures similarity between $\overline{H}_s$ and the true spatial gene expression $H_s$ by optimizing a contrastive loss function. The loss maximizes similarities for positive pairs (spatially adjacent spots) and minimizes similarities for negative pairs (spatially non-adjacent spots), preserving the raw topological structure of spatial locations.

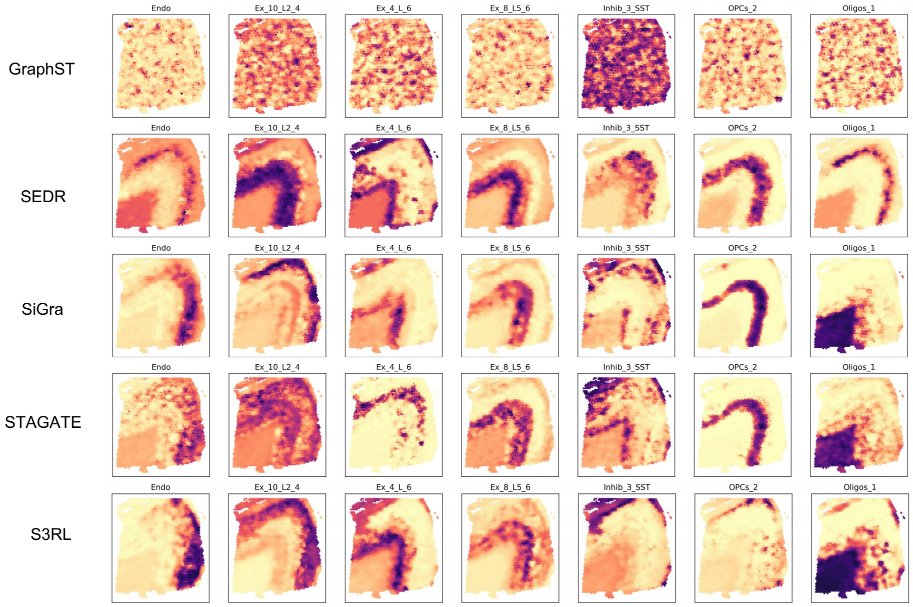

**Fig. S6: Visualization of deconvolution results across methods on DLPFC slice 151673.** Comparison of cell-type proportion maps generated by different reconstruction-capable methods (GraphST, SEDR, SiGra, STAGATE, and our proposed S3RL) under the same deconvolution framework. All models share the same contrastive deconvolution strategy as proposed in this work. Despite the lack of precise quantitative evaluation, visual inspection shows that S3RL yields the clearest spatial boundaries and the most coherent cell-type distributions, especially for major cell types such as Ex (excitatory neurons), Oligos (oligodendrocytes), and Endo (endothelial cells). These results further demonstrate the strong reconstruction quality and spatial consistency achieved by S3RL.

#### 2 Latent Space Clustering Results on NanoString dataset

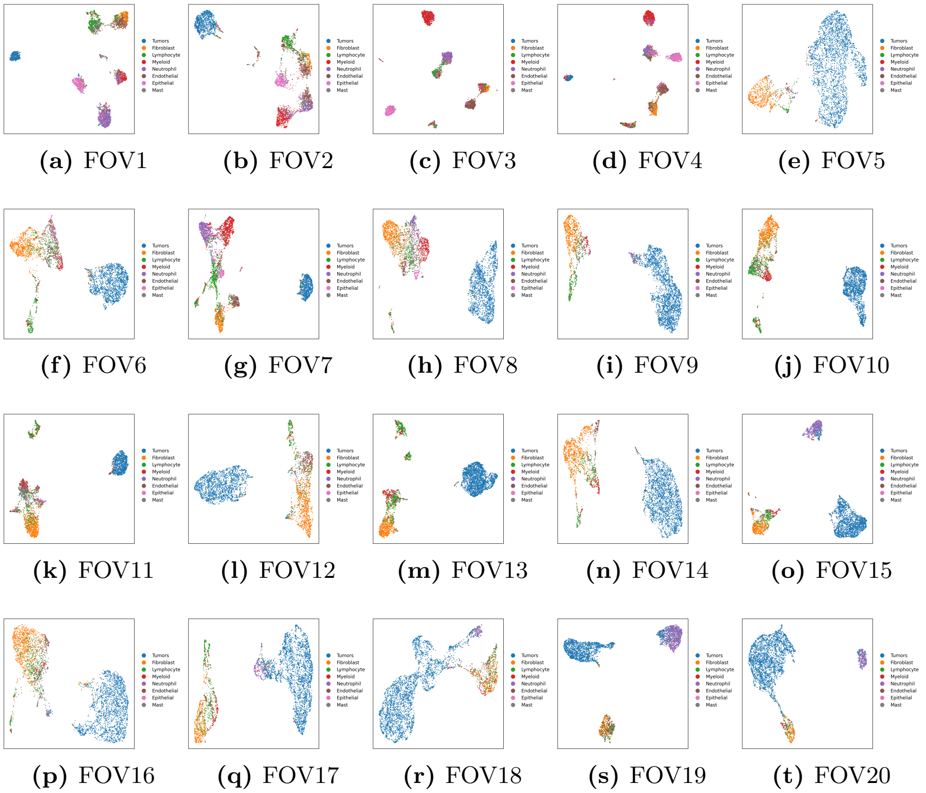

**(a)** FOV1 **(b)** FOV2 **(c)** FOV3 **(d)** FOV4 **(e)** FOV5

**(f)** FOV6 **(g)** FOV7 **(h)** FOV8 **(i)** FOV9 **(j)** FOV10

**(k)** FOV11 **(l)** FOV12 **(m)** FOV13 **(n)** FOV14 **(o)** FOV15

**(p)** FOV16 **(q)** FOV17 **(r)** FOV18 **(s)** FOV19 **(t)** FOV20

**Fig. S7: Clustering results in the latent space for 20 lung cancer slices from the Nanostring dataset using S3RL.** Each subplot represents the latent space embedding of one slice, where spots are colored according to their predicted clusters. The clustering results highlight the spatial organization of different cell populations.

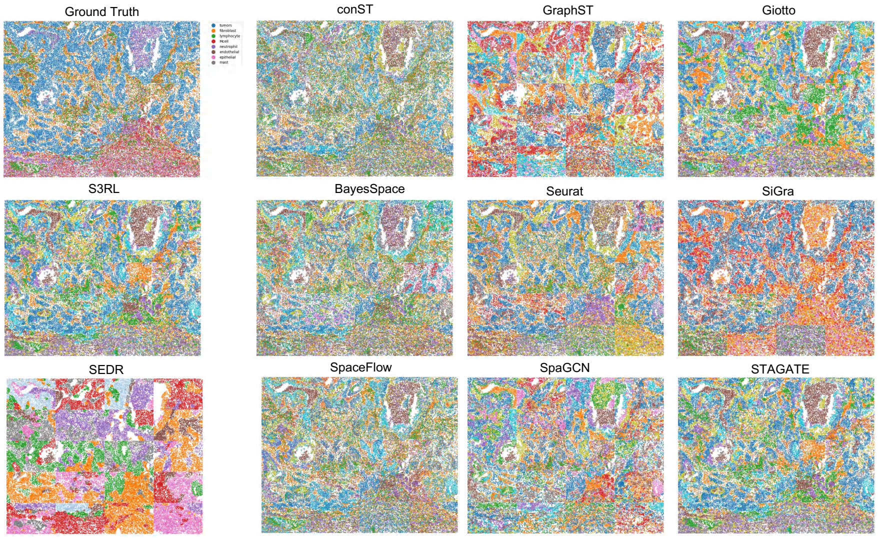

**Fig. S8: Comparison of clustering results across 20 Nanostring lung cancer slices using different methods.** The first image (top-left) represents the ground truth annotations, while the remaining images depict clustering results obtained from various spatial transcriptomics analysis frameworks, including conST, GraphST, Giotto, S3RL, BayesSpace, Seurat, SiGra, SpaceFlow, SpaGCN, and STAGATE. Our proposed S3RL method demonstrates improved spatial consistency and boundary precision in capturing cellular heterogeneity across lung tissue samples.

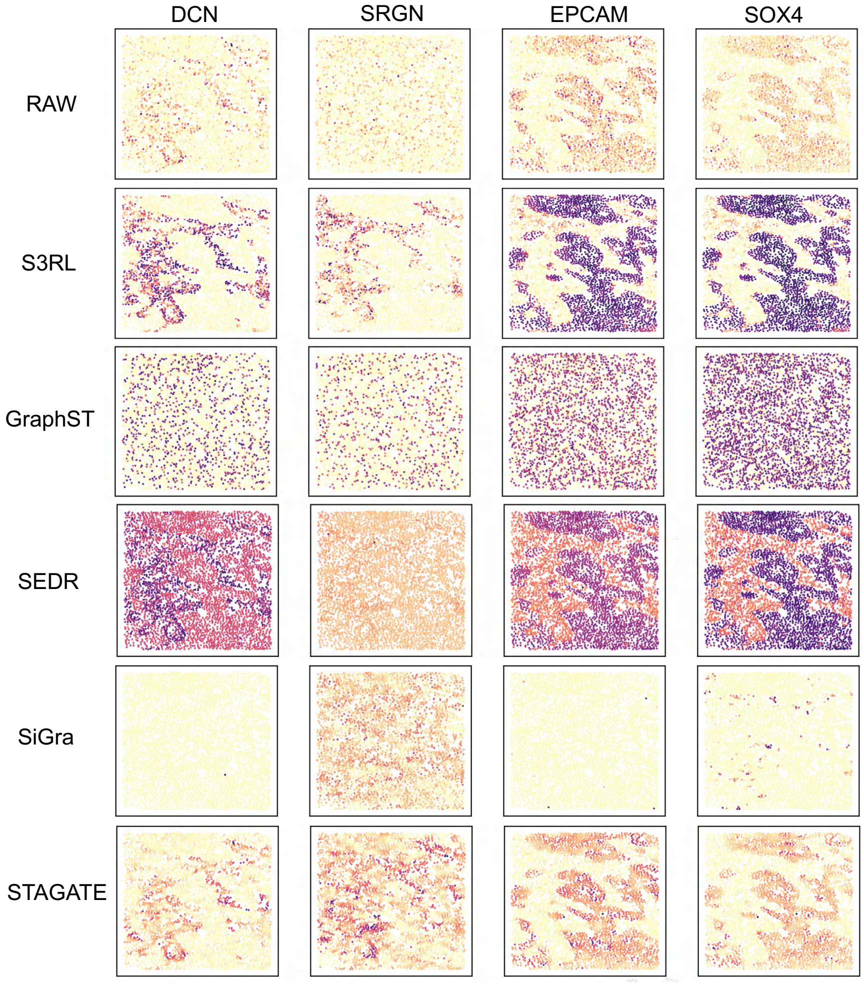

**Fig. S9: Visualization of marker gene reconstruction performance across generative spatial transcriptomics models.** Comparison of reconstructed spatial expression patterns for four key marker genes (*DCN*, *SRGN*, *EPCAM*, and *SOX4*) across different methods, including the raw ST data (top row), and the reconstructed outputs of S3RL, GraphST, SEDR, SiGra, and STAGATE. All methods were evaluated under the same reconstruction framework using their model-generated latent outputs. The S3RL reconstruction shows the clearest spatial boundaries and expression contrast, especially for *SRGN*, where it achieves a more distinguishable region separation compared to other methods. These results demonstrate the effectiveness of S3RL in capturing biologically relevant gene expression patterns during spatial reconstruction.

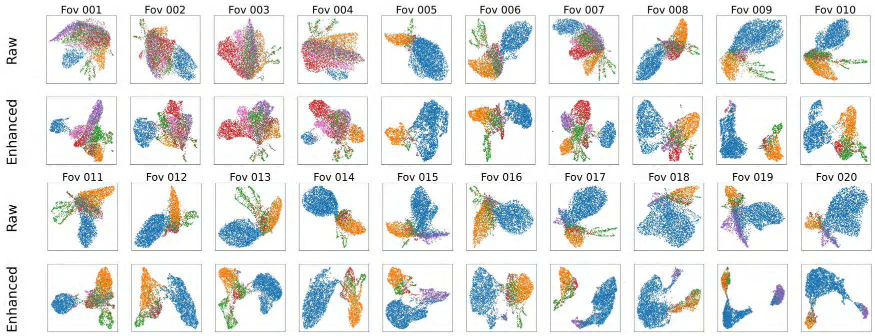

**Fig. S10: Comparison of the latent space clustering performances in raw and enhanced clustering for Nanostring lung cancer slice data.** The first and third rows show unprocessed data, while the second and fourth depict S3RL-enhanced embeddings. These enhanced embeddings illustrate cluster separability improved spatially and with cell-type distinction based on more appropriate clustering within the tumor microenvironment. Again, S3RL shows promise in spatial transcriptomics studies.

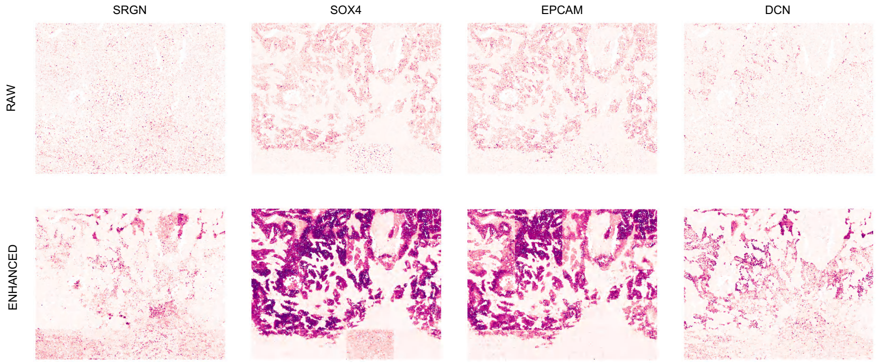

**Fig. S11: Comparison of marker gene expression in raw and enhanced data using the S3RL in Nanostring lung cancer slice data**. The top row is raw data, and the bottom row is enhanced data. The four selected markers (SRGN, SOX4, EPCAM, and DCN) are critical in lung cancer: SRGN is influenced by the tumor microenvironment modulation, SOX4 is a transcription factor that regulates epithelial-mesenchymal transition (EMT), EPCAM is an epithelial cancer marker associated with metastasis, and DCN (Decorin) is involved in ECM remodeling. The data enhancement clearly shows spatial expression patterns more consistent with expected tissue structure, underscoring S3RL's utility for resolution and biological interpretability.

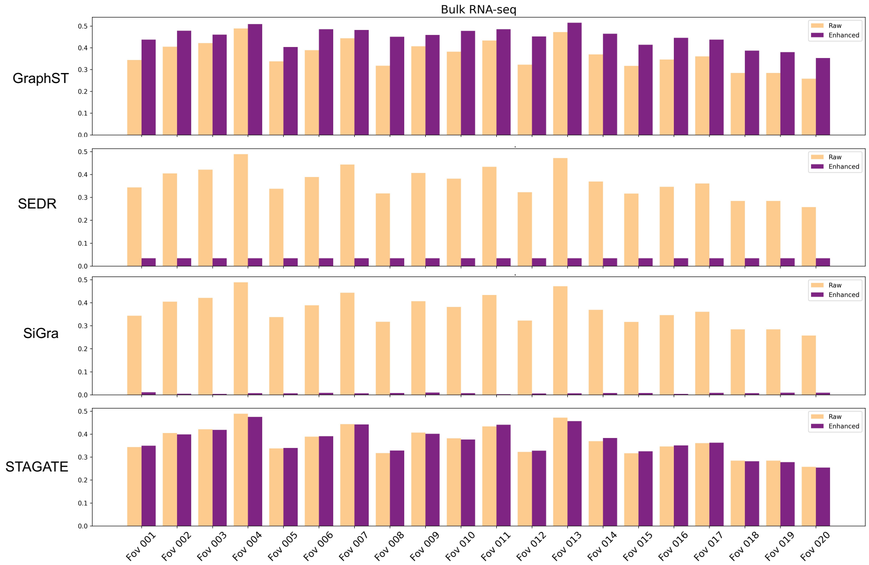

**Fig. S12: Comparison of Pearson correlation between bulk RNA-seq (TCGA-LUAD) and raw/enhanced spatial transcriptomic data across 20 Nanostring lung slices of different methods.** Bar plots showing the Pearson correlation between bulk RNA-seq (TCGA-LUAD) and spatial transcriptomic data (raw vs. enhanced) across 20 Nanostring lung slices for four reconstruction-capable methods: GraphST, SEDR, SiGra, and STAGATE.

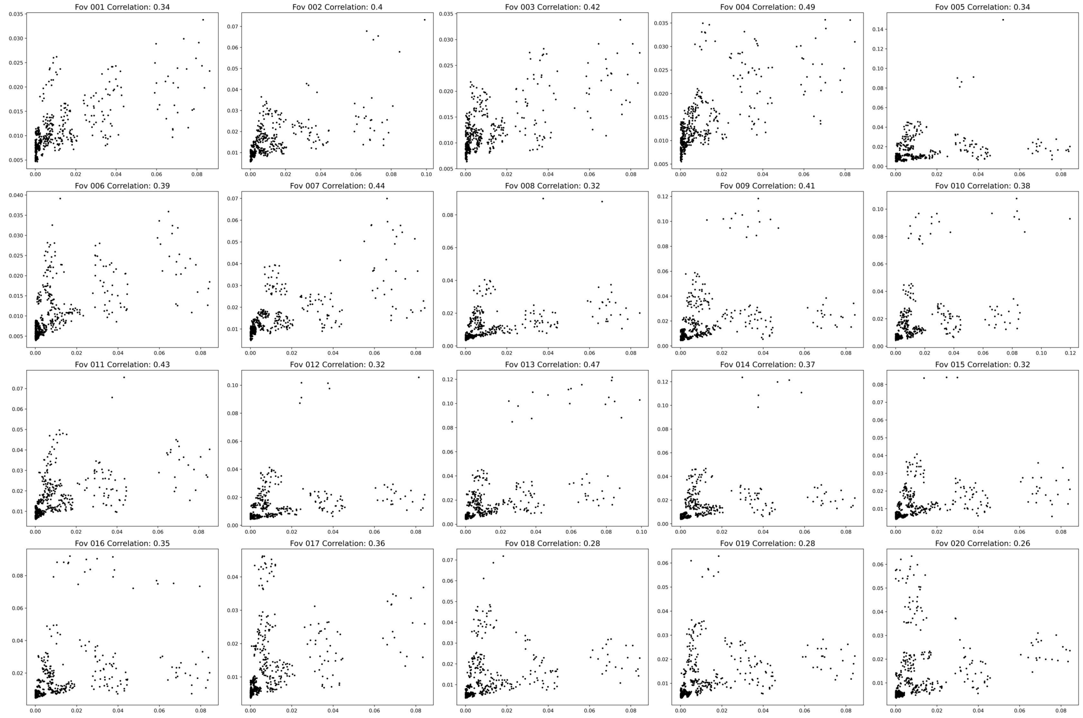

**Fig. S13: Correlation analysis between bulk RNA-seq and raw spatial transcriptomics data in the Nanostring dataset.** Each scatter plot corresponds to one tissue slice (FOV), where each point represents a gene. The x-axis and y-axis denote gene expression levels in the bulk RNA-seq and raw spatial transcriptomics data, respectively. The plots illustrate the correlation of expression patterns between bulk and spatial data under raw conditions, showing that S3RL enhancement improves consistency across most slices.

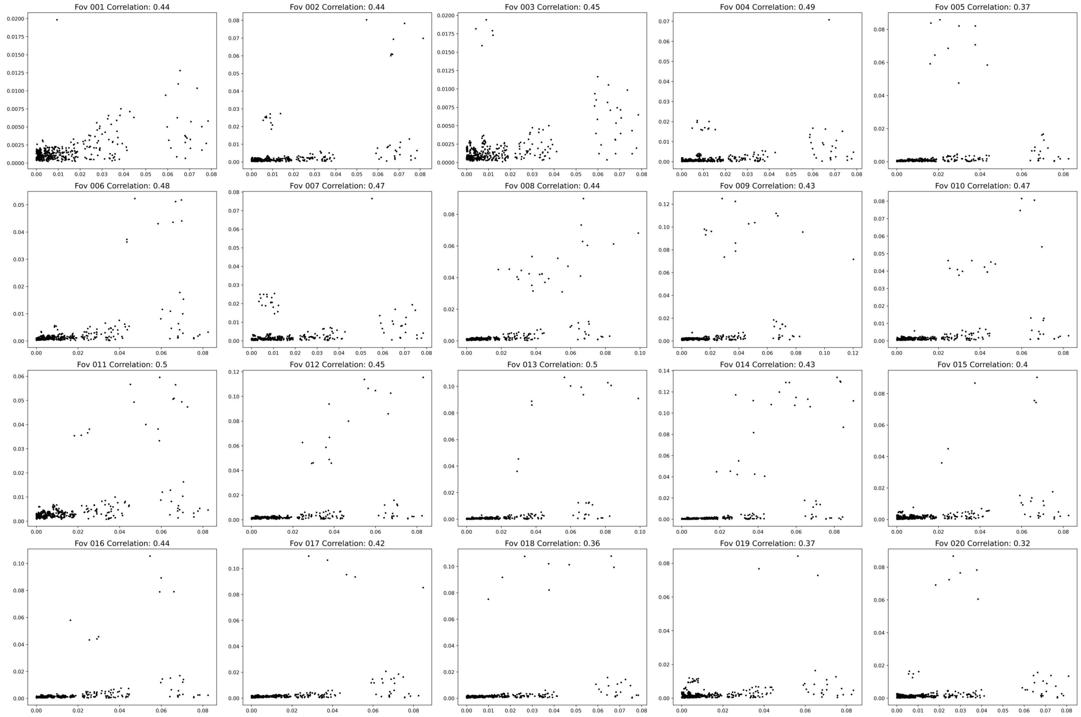

**Fig. S14: Correlation analysis between bulk RNA-seq and enhanced spatial transcriptomics data in the Nanostring dataset.** Each scatter plot corresponds to one tissue slice (FOV), where each point represents a gene. The x-axis and y-axis denote gene expression levels in the bulk RNA-seq and enhanced spatial transcriptomics data, respectively. The plots illustrate the correlation of expression patterns between bulk and spatial data under enhanced conditions, showing that S3RL enhancement improves consistency across most slices.

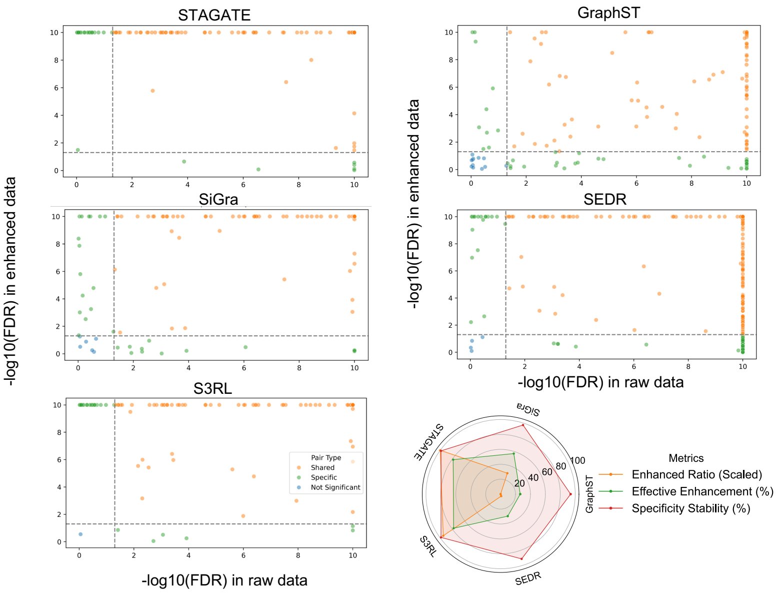

**Fig. S15: Comparison of ligand-receptor (L-R) pair significance across different enhancement methods.** Scatter plots show the significance level of L-R pairs based on $-\log_{10}(\text{FDR})$ in raw data (x-axis) and enhanced data (y-axis) for five methods: STAGATE, GraphST, SiGra, SEDR, and S3RL. Each dot represents an L-R pair, colored by its classification: shared (orange), enhanced-specific (green), or not significant (blue). The radar plot summarizes three evaluation metrics across methods: Enhanced Ratio (scaled), Effective Enhancement (%), and Specificity Stability (%).

### 3 Spatial Clustering Results on Mouse Brain Anterior and Human Breast Cancer Datasets

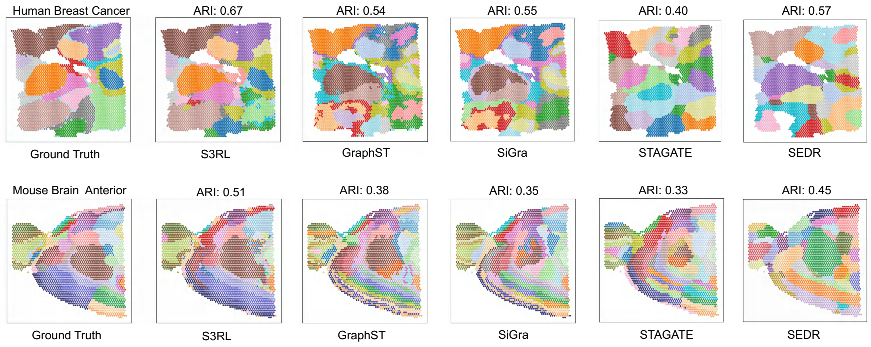

**Fig. S16: Clustering comparison on Mouse Brain Anterior and Human Breast Cancer datasets.** The spatial clustering results obtained by different methods (GraphST, SEDR, SiGra, STAGATE, and S3RL) on two representative datasets (10X Visium spatial transcriptomics datasets): Mouse Brain Anterior (top) and Human Breast Cancer (bottom). Ground truth annotations are provided as references. Adjusted Rand Index (ARI) values are reported above each result to quantify clustering accuracy. The results demonstrate that S3RL achieves higher ARI scores and better boundary alignment compared to other methods, providing more accurate spatial domain segmentation in both neural and tumor tissues.

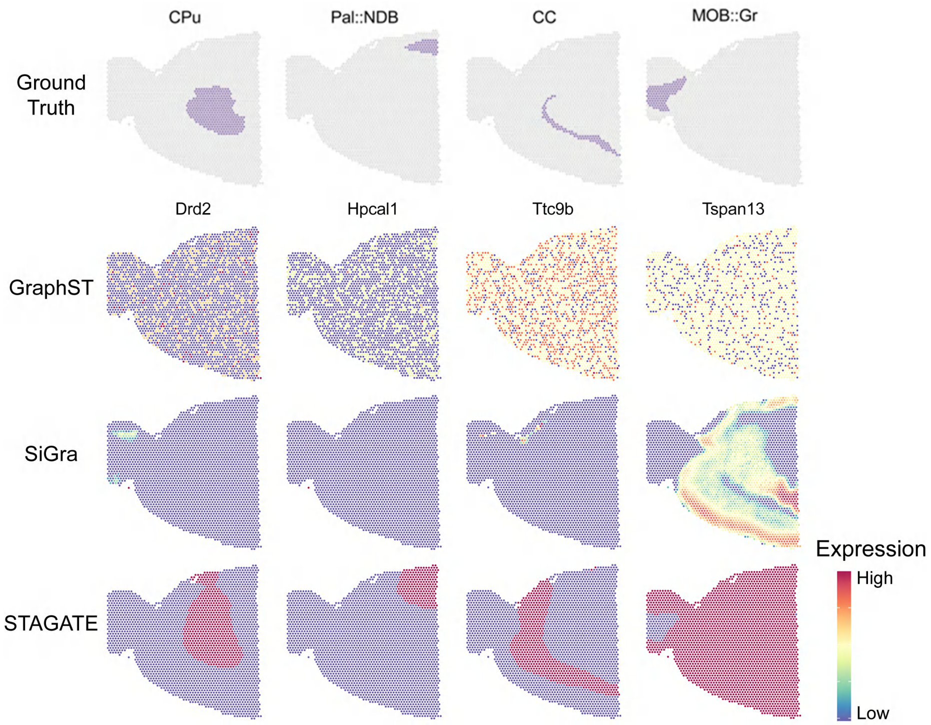

**Fig. S17:** Comparison of reconstructed marker gene expression patterns by baseline methods on the Mouse Brain Anterior dataset. The top row shows the ground truth cell-type annotations for each spatial region, while each subsequent row presents the spatial expression patterns of selected marker genes reconstructed by baseline models, including STAGATE, GraphST, and SEDR. Although certain patterns are partially recovered, the reconstructions often appear spatially diffused or lack boundary specificity, making it difficult to clearly delineate anatomical regions. This highlights the limitations of current generative methods in recovering fine-grained spatial gene expression in complex neural tissues.

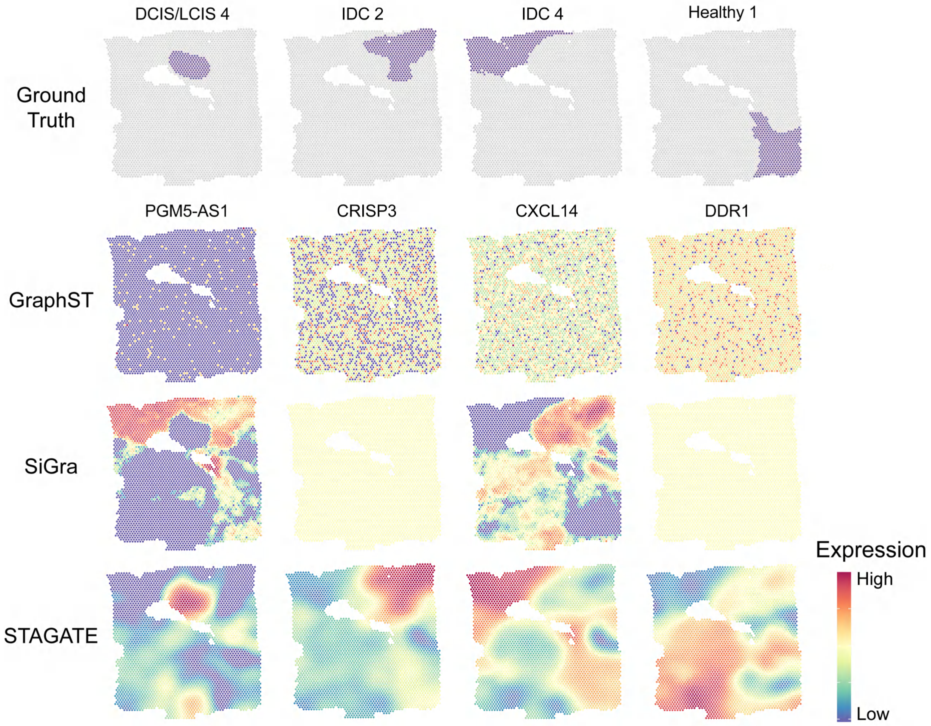

**Fig. S18: Reconstruction of marker gene expression in human breast cancer tissue by different baseline methods.** The top row displays ground truth annotations of spatial domains based on expert interpretation. The rows below show the reconstructed spatial expression patterns of key marker genes by different generative models. While some spatial structures are partially recovered, most methods exhibit diffuse or blurred boundaries, making it difficult to distinguish fine-grained tumor microenvironmental regions. This highlights the current limitations of generative frameworks in accurately capturing spatial heterogeneity in complex cancer tissues.
